# Translation inhibition efficacy does not determine the *Plasmodium berghei* liver stage antiplasmodial efficacy of protein synthesis inhibitors

**DOI:** 10.1101/2023.12.07.570699

**Authors:** James L. McLellan, Kirsten K. Hanson

## Abstract

Protein synthesis is a core cellular process, necessary throughout the complex lifecycle of *Plasmodium* parasites, thus specific translation inhibitors would be a valuable class of antimalarial drugs, capable of both treating symptomatic infections in the blood and providing chemoprotection by targeting the initial parasite population in the liver, preventing both human disease and parasite transmission back to the mosquito host. As increasing numbers of antiplasmodial compounds are identified that converge mechanistically at inhibition of cytoplasmic translation, regardless of molecular target or mechanism, it would be useful to gain deeper understanding of how their effectiveness as liver stage translation inhibitors relates to their chemoprotective potential. Here, we probed that relationship using the *P. berghei*-HepG2 liver stage infection model. Using o-propargyl puromycin-based labeling of the nascent proteome in *P. berghei-*infected HepG2 monolayers coupled with automated confocal feedback microscopy to generate unbiased, single parasite image sets of *P. berghei* liver stage translation, we determined translation inhibition EC_50s_ for five compounds, encompassing parasite-specific aminoacyl tRNA synthetase inhibitors, compounds targeting the ribosome in both host and parasite, as well as DDD107498, which targets *Plasmodium* eEF2, and is a leading antimalarial candidate compound being clinically developed as cabamiquine. Compounds were then tested at equivalent effective concentrations to compare the parasite response to, and recovery from, a brief period of translation inhibition in early schizogony, with parasites followed up to 120 hours post-infection to assess liver stage antiplasmodial effects of the treatment. Our data conclusively show that translation inhibition efficacy *per se* does not determine a translation inhibitor’s antiplasmodial efficacy. DDD107498 was the least effective translation inhibitor, yet exerted the strongest antimalarial effects at both 5x- and 10x EC_50_ concentrations. We show compound-specific heterogeneity in single parasite and population responses to translation inhibitor treatment, with no single metric strongly correlated to release of hepatic merozoites for all compound, demonstrate that DDD107498 is capable of exerting antiplasmodial effects on translationally arrested liver stage parasites, and uncover unexpected growth dynamics during the liver stage. Our results demonstrate that translation inhibition efficacy cannot function as a proxy for antiplasmodial effectiveness, and highlight the importance of exploring the ultimate, as well as proximate, mechanisms of action of these compounds on liver stage parasites.

## Introduction

*Plasmodium* parasite resistance to antimalarial drugs is a serious problem throughout endemic regions and underscores the need for new antimalarial drugs that work against novel drug targets (*1*). In addition to killing asexual blood stage parasites, which are solely responsible for the disease, malaria, new compounds capable of killing *Plasmodium* parasites at multiple life cycle stages, including the obligate liver stage of development, are also greatly needed (*2*).

Recently, several compounds were identified with the ability to selectively target parts of the *Plasmodium* cytoplasmic translation machinery, including diverse amino acyl-tRNA synthetase enzymes (reviewed in (*3, 4*)), and eukaryotic elongation factor 2 (eEF2) (*5*), most of which display multistage antiplasmodial activity. The eEF2 inhibitor, DDD107498 (also known as M-5717, and currently developed as cabamiquine), has successfully completed phase I trials, with demonstrated efficacy against both liver stage and blood stage *P. falciparum* infection in human volunteers (*6, 7*). The promise of compounds targeting *Plasmodium* protein synthesis has led to increased efforts to identify additional novel compounds and rational drug targets in the *Plasmodium* cytoplasmic translation apparatus (*3, 4, 8–11*). As an increasing number of *Plasmodium-*specific translation inhibitors emerge from these collective efforts, it would be useful to be able to identify compounds with the most desirable activity profiles. Regardless of their molecular target, these compounds converge mechanistically, in that all target the process of cytoplasmic translation. This calls into question the relationship between mechanistic efficacy, how effective such a compound is at inhibiting translation *in cellulo*, and antiplasmodial efficacy, its ability to kill parasites.

Though most mechanistic antimalarial work has been confined to the asexual blood stage (ABS), translation inhibitors largely have multistage antimalarial activity, and the demand for protein synthesis during liver stage (LS) development far exceeds that of the ABS; the LS is essentially an amplification step, where a single invading sporozoite will grow and replicate inside a hepatocyte to generates thousands of progeny capable of initiating the blood stage of infection (*12*). During the ABS, a single merozoite invasion event will generate ∼10-32 new erythrocytic merozoite progeny in a single 48 h (hour) replication cycle in *P. falciparum* (e.g. (*13, 14*)), while the rodent model parasite *P. berghei* will generate on average ∼12 erythrocytic merozoites in a 24 h cycle (*15*). In contrast, a single *P. berghei* sporozoite hepatocyte invasion event can generate 1500 - 8000 merozoite progeny during the 2-3 days of LS development, while a single *P. falciparum* LS schizont generates 30-40,000 hepatic merozoites during a 6 day growth period ((*16*), and references therein). Since the biosynthetic output needed to support LS growth and hepatic merozoite production is so much greater than that required for erythrocytic merozoites production, replicating LS parasites might be particularly sensitive to translation inhibition.

Previously, we developed a single-cell quantitative translation assay for *P. berghei* liver stage parasites (*17*), also known as exoerythrocytic forms (EEFs), using o-propargyl puromycin (OPP) (*18*), to label the LS nascent proteome *in cellulo*, and showed that liver stage translation inhibition efficacy varied between known translation inhibitors when tested at high, presumably saturating, concentrations. Our estimated translation inhibition EC_50_ for DDD107498 in *P. berghei* LS was 13.4 nM (*17*), substantially higher than the EC_50_s for antiplasmodial activity reported against *P. berghei* and *P. yoelii* liver stages and *P. falciparum* asexual blood stages (ABS), which were 1.65 nM, 0.97 nM, and 1 nM, respectively (*5, 19*).

DDD107498 was also a less efficacious inhibitor of *P. berghei* LS translation at a high, presumably saturating concentration, than others tested (*17*). These differences are somewhat surprising, given the extremely strong genetic evidence that *P. falciparum* ABS parasites evolve resistance to DDD107498 *in vitro*, in mouse models, and in humans via mutations in the *P. falciparum* eukaryotic elongation factor 2(eEF2) gene (*5, 6, 20, 21*), encoding a highly conserved GTPase required to catalyze translocation during the translation elongation process. Taken together, these differences led us to question whether a translation inhibitor’s antiplasmodial efficacy — defined here, with respect to the liver stage, as the ability to prevent the formation and release of hepatic merozoites— is driven by the translation inhibition efficacy of the compound. We thus decided to use the *P. berghei*-HepG2 infection model to probe the relationship between translation inhibition efficacy and antiplasmodial efficacy by examining the effect of a brief period of translation inhibition during LS schizogony on parasite protein synthesis, growth, hepatic merozoite production and merosome / detached cell release comparing five mechanistically distinct translation inhibitors tested at equivalent effective concentrations.

## Results

### Translation inhibitor characteristics and potency determination

As our goal was to study the LS parasite response to a brief period of translation inhibition when the translational output was high, we first used OPP-labeling of the *P. berghei* LS nascent proteome to confirm our previous result that parasites at 28 hours post infection (hpi) have a greater translational intensity than those at 48 hpi (*17*). As the average translational intensity at 28 hpi was more than double that at 48 hpi (Figure S1A), we elected to use a 4 hour (h) treatment window from 24-28 hpi, with nascent proteome labeling from 27.5-28 hpi in our experiments. We selected five compounds representing a diverse collection of molecular targets, modes/mechanisms of action, parasite vs. host selectivity, and potencies.

Four of these compounds were used in our previous work to develop the LS translation assay: anisomycin, bruceantin, DDD107498, and MMV019266. Anisomycin is a pan-eukaryotic elongation inhibitor that binds the ribosomal A-site (*22*), and has similar activity against *P. berghei* LS translation and that of HepG2 cells (*17*). Despite this dual activity and high translation inhibition efficacy, a 24-28hpi treatment with a high concentration of anisomycin was completely reversible, in terms of parasite translation levels, 20 h after compound washout (*17*). Bruceantin, a translation elongation inhibitor that binds the ribosomal A-site and only efficiently inhibits monosomes (*23, 24*) was both the most potent and most efficacious inhibitor *of P. berghei* LS translation, and also induced the most variable single parasite outcomes at concentrations evoking submaximal effects (*17*); four hours of treatment with 3.7 nM bruceantin led to extremely heterogeneous responses in individual parasites, with ∼83% of the sampled population translationally inhibited, while the remaining ∼17% appeared unaffected (Figure S1B). In contrast, parasites treated with the aforementioned DDD107498 maintained a normal population distribution at concentrations evoking both submaximal and maximal effects (Figure S1C). MMV019266, a multistage active Malaria Box compound (*25*), which was identified in our previous study as a *Plasmodium*-specific translation inhibitor (*17*), is likely to target the cytoplasmic isoleucyl-tRNA synthetase, as has been demonstrated for several structurally related thienopyrimidines (*26*). Using a parasite reduction ratio (PRR) assay (*27*), 10xEC_50_ MMV019266 was shown to kill the majority of *P. falciparum* ABS parasites after 24h of treatment (*26*), a rapid rate-of-kill similar to that of the fast-acting reference antimalarial artesunate. This rapid ABS rate-of-kill of MMV019266 is in marked contrast to that of DDD107498, which does not reduce the viable parasite population at all after a 24 h 10xEC_50_ treatment (*5*), and has a PRR similar to that of atovaquone, the slowest acting reference antimalarial in the assay. We also chose to test another recently identified parasite-specific aminoacyl-tRNA synthetase (aaRS) inhibitor, LysRS-IN-2, identified in a target-based screen of compounds capable of inhibiting *P. falciparum* cytoplasmic lysyl-RS activity (*28*); LysRS-IN-2 has an ABS speed-of-kill even slower than that of atovaquone (*28*). Using a competitive OPP (co-OPP) assay (*17*), we demonstrate that LysRS-IN-2 is a specific, and likely direct, inhibitor of *P. berghei* LS translation (Figure S1D).

We first determined the translation inhibition potency of each selected inhibitor, tested in 8- or 10-point, 3-fold serial dilution, with 3.5 hours of pre-treatment followed by 30 minutes of o-propargyl puromycin (OPP) labeling in the presence of the compound of interest (Figure 1A-B, see Table S1 for EC_50_, 95%CI, and R square). As we previously demonstrated (*17*), the pan-eukaryotic translation inhibitors anisomycin and bruceantin are active against both *P. berghei* and HepG2 translation, but display differences in potency and selectivity. Anisomycin is nearly equipotent against *P. berghei* EEFs and HepG2 cells with EC_50_ values of 262 nM and 198 nM respectively (Figure 1A). Both host and parasite show a small concentration-dependent increase in translation initially before switching to translation inhibition, with the entire population of single parasites shifted (Figure 1A-B). Bruceantin is more active against *P. berghei*, with an EC_50_ of 3.72 nM compared to 13.7 nM in HepG2 (Figure 1A). At the single parasite level (Figure 1B), bruceantin appears to be the only compound tested for which a concentration that evoked submaximal translation inhibition shows the parasite population effectively split into a group of strongly responding parasites, the majority, and those that show little if any translation inhibition. Of the three parasite specific translation inhibitors, DDD107498 was the most potent, with a translation inhibition EC_50_ of 11.3 nM, the MMV019266 EC_50_ was 456 nM, and the least potent compound tested was LysRS-IN-2, with an EC_50_ of 6630 nM (Figure 1A).

**Figure 1.**
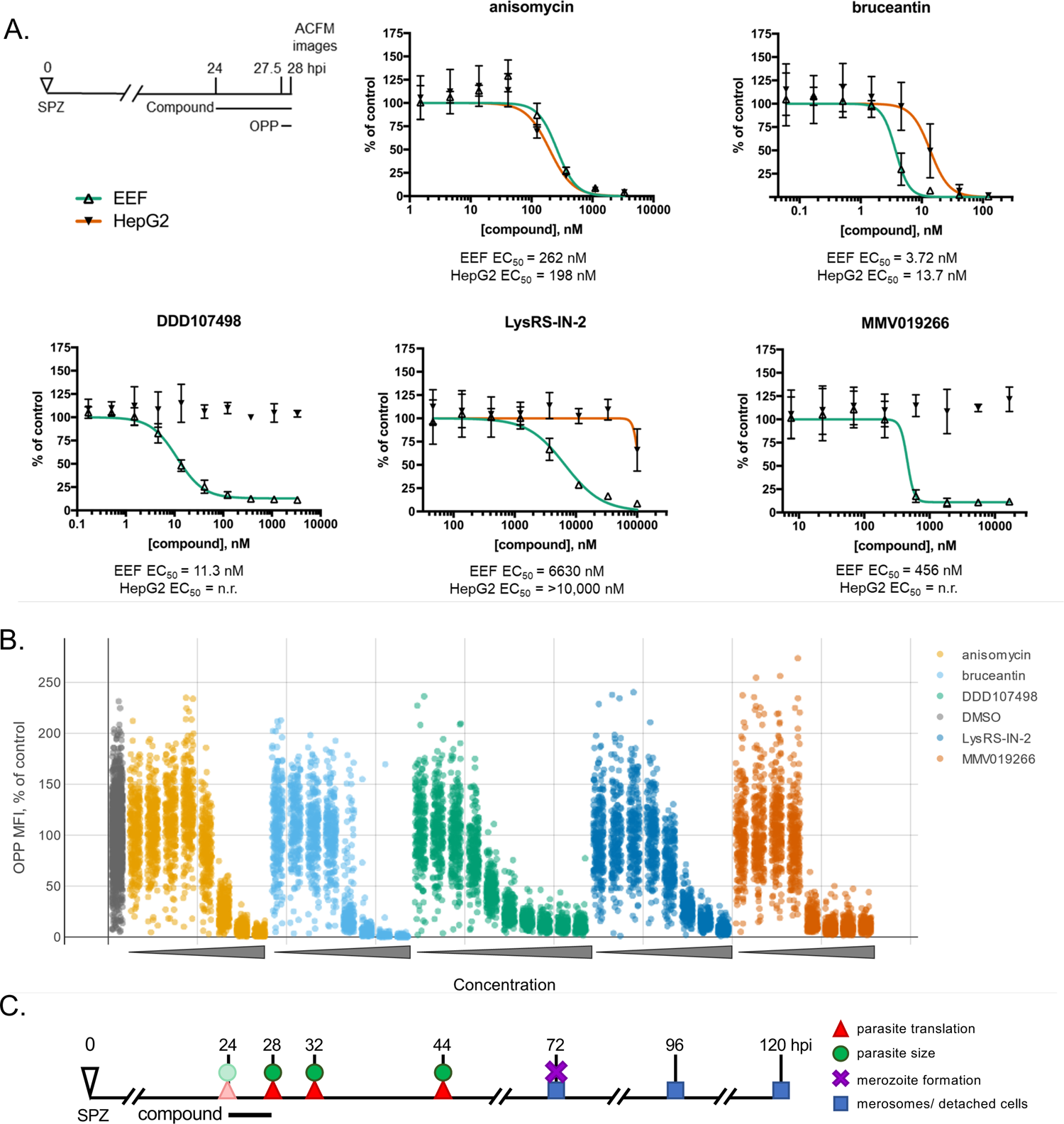
Translation inhibition potency determination and experimental overview. Translation inhibition was quantified in *P. berghei* EEFs (A-B) and matching in-image HepG2 (A) following acute pre-treatment from 24 - 28 hpi, as shown in the schematic. Compounds were tested in 8 or 10 point, 3-fold serial dilution. (A) Concentration response curves. Single data points represent the mean translation at each concentration, normalized to DMSO controls, and errors bars show standard deviation from n=3 independent experiments. (B) Single LS parasite concentration response. For each compound, concentrations plotted moving from lowest (left) to highest (right) as indicated by the triangles. Top concentrations tested were 3,333 nM for anisomycin and DDD107498, 123 nM for bruceantin, 16,667 nM for MMV01266, and 100,000 nM for LysRS-IN-2. N=3 independent experiments combined, each dot represents mean translation intensity (OPP-MFI) in a single parasite normalized to DMSO controls, n=10,461EEFs total. The full concentration-response dataset can be explored via interactive dashboards in in our KNIME hub workflow: https://hub.knime.com/-/spaces/-/~TZCrKvv3sbJwM_xP/current-state/. (C) Experimental setup to probe the effects of acute translation inhibition and the relationship between translation inhibition efficacy and antiplasmodial efficacy with inhibitors tested at equivalent effective concentrations.

Canonical LS compound activity assays, with treatment throughout LS development until a 48 hpi readout of parasite biomass, demonstrated that DDD107498 was ∼9.4 fold less potent in inhibiting LS translation than in inhibiting LS growth, while LysRS-IN-2 and MMV019266 were ∼3.6- and ∼2.5 fold less potent, respectively (Table S1). DDD107498 had the shallowest slope of the compounds tested, while MMV019266 had a markedly steeper slope than the other 4 compounds (Figure 1A-B). Even though saturating effects on translation were achieved with both DDD107498 and MMV019266, they were noticeably less efficacious than anisomycin and bruceantin, where parasite translation is nearly undetectable (Figure 1A).

To probe the relationship between translation inhibition efficacy and antiplasmodial efficacy in the LS, we designed an experiment to quantify parasite translation and size at the end of 4 treatment with 5x and 10x EC_50_ concentrations of each compound, then, after compound washout, monitoring translation and EEF size, along with late liver stage maturation, merozoite formation, and merosome/detached cell release in experimentally matched samples (Figure 1B). Additionally, DDD107498 was also tested at a lower concentration corresponding to 10x EC_50_ for LS biomass reduction; as this EC_50_ value falls between 1 and 2 nM, we decided to round up, and use a 20 nM concentration, which corresponds to 1.8x our calculated translation inhibition EC_50_. At 24 hpi, pre-treatment controls were OPP-labeled and fixed to determine a baseline for parasite translational output and size, and the remaining samples were compound treated. At 28 hpi, a subset of the treated samples was OPP-labeled and fixed to quantify translation inhibition and parasite size at the end of the 4 h treatment. The remaining treated samples were subjected to a stringent compound washout protocol, where the wash volume and total number of media exchanges was calculated to ensure a ≥ 4-log reduction in compound concentration per well, e.g. the concentration in the 10x EC_50_ treatments after washout was reduced to ≤ 0.001x EC_50_. After the washout, parasite recovery was assessed via both translational intensity and size at 32 hpi (4 hours after washout) and 44 hpi (16 hours after washout). After processing, individual parasites from the 24-, 28-, 32-, and 44 hpi samples were imaged via automated confocal feedback microscopy (*17, 29*). At 72 hpi, LS parasite maturation and hepatic merozoite formation were assessed by quantifying the percentage of monolayer EEFs expressing merozoite surface protein 1 (MSP1) and apical membrane antigen 1 (AMA1) in immunolabeled monolayers; we collected the culture medium from the same wells to quantify merozoite-filled, detached HepG2 cells, the presentation of merosome formation in the *P. berghei*-HepG2 infection model (*30*). In parallel, independent samples were set up for repeat merosome collection every 24 hours at 72-, 96- and 120 hpi, with complete medium replacement after each collection. Merosome release in the *P. berghei*-HepG2 infection model is considered largely completed by 65 hpi (*30, 31*), and the continually replicating HepG2 cells begin to overgrow the monolayer, but we hoped that we could separate potential compound-induced developmental delays, or cytostatic effects, from those that likely represent parasite killing, or cytotoxic effects.

### Translation inhibition efficacy varies between compounds

Translation inhibition efficacy was quantified in image sets generated by automated confocal feedback microscopy (ACFM) in both *P. berghei* liver stage parasites (Figure 2A & see Figure S2A for data reproducibility between experiments) and the in-image HepG2 cells (Figure S4) at 28 hpi after 4 h of treatment. As expected, parasite translation was significantly inhibited for all compound concentrations tested compared to the DMSO control (one-way ANOVA, p<0.0001). Additionally, substantial differences in the ability to inhibit parasite translation were seen between the compounds at both the 5x- and 10x EC_50_ concentrations (one-way ANOVA; 5x EC_50_ p<0.0001, 10x EC_50_ p=0.002; for Tukey’s multiple comparisons of experimental means, see Table S2). All compounds displayed greater inhibitory activity at 10x EC_50_ than 5x EC_50_, except for MMV019266 and bruceantin where the two concentrations induced essentially equivalent effects (Figure 2A & S2A, Table S2A). MMV019266 and LysRS-IN-2 were the most effective of the parasite-specific inhibitors at 5x- and 10x EC_50_, respectively, while DDD107498 was the least effective of the parasite-specific translation inhibitors at both effective concentrations (Figure 2B & S2A). MMV019266 and Lys-IN-2 caused >90% translation inhibition at both concentrations, but DDD107498 never reached 90% at either 5x- or 10x EC_50_ and resulted in only 64% translation inhibition at 1.8x EC_50_ (Table S2). The two pan-eukaryotic inhibitors bruceantin and anisomycin were both highly efficacious; on average, the percent LS translation inhibition by bruceantin was >99% in both 5x- and 10x EC_50_ concentrations, while anisomycin-induced translation inhibition was approximately 97% and 99% at 5x- and 10x EC_50_, respectively (Figure 2A, S2A, and Table S2).

**Figure 2.**
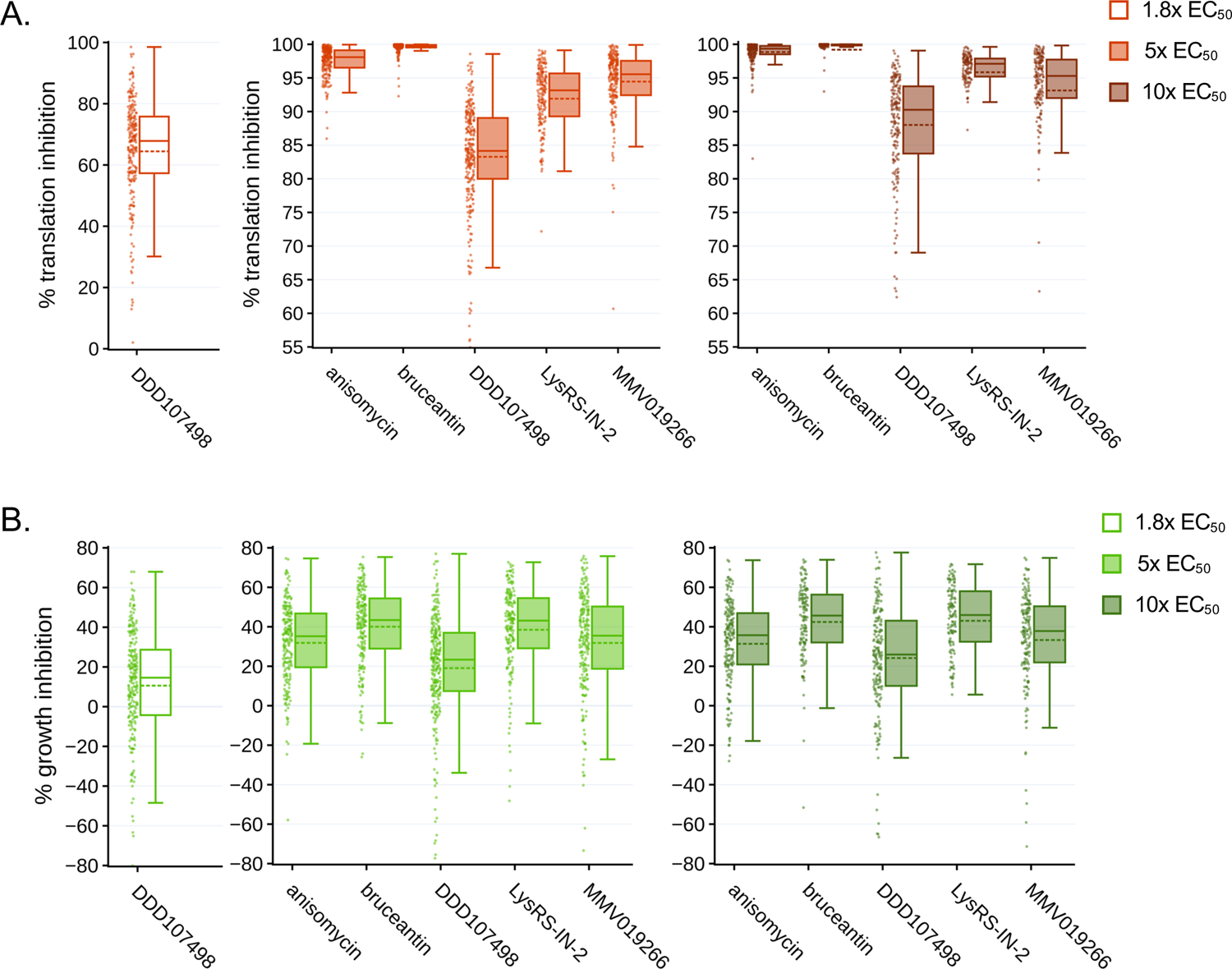
Differential effects of translation inhibitors tested at equivalent effective concentrations. Translation inhibition (A) and growth inhibition (B) at 28 hpi in *P. berghei* LSs, following 4 h acute pre-treatments. Individual data points represent single parasites normalized to in-plate DMSO controls set to 0. Boxplots show combined data from all 3 independent experiments, with dotted lines reporting treatment means. Statistical significance was assessed using a one-way ANOVA with Tukey’s multiple comparisons testing, reported in table S2. To facilitate an easier comparison between treatments, plots were truncated at 55% (A) and −80% (B), removing a total of 8 and 12 individual outlier EEFs respectively. The full dataset can be explored via interactive dashboards in in our KNIME hub workflow: https://hub.knime.com/-/spaces/-/~EcnvMwYtqylu2reV/current-state/.

We also quantified parasite size after acute translation inhibition (Figure 2B and see Figure S2B for data reproducibility between experiments). Compared to DMSO controls, significant reductions in parasite growth were seen for all compound concentrations tested (one way ANOVA; p<0.0001), and significant differences in the growth inhibition were seen between compounds at 5x and 10x EC_50_ concentrations (one way ANOVA; 5x EC_50_, p=0.0006, 10x EC_50_ p=0.0028; for Tukey’s multiple comparisons of experimental means see Table S2). Due to the causal relationship between protein synthesis and cell growth, we hypothesized that the rank order of compounds for LS growth inhibition at each multiple of EC_50_ would be identical to that of LS translation inhibition, but while the most and least effective translation inhibitors correspondingly caused the most and least parasite growth inhibition at both 5x- and 10x EC_50_, the other three compounds did not rank equivalently (Figure 2B and see Table S2 for mean values per treatment). Parasite growth was most strongly inhibited in bruceantin and LysRS-IN-2 treatments, where EEF growth inhibition was approximately 39% and 43% of DMSO controls in the 5x- and 10x EC_50_ treatments, respectively, despite bruceantin inducing a significantly stronger effect on translation than LysRS-IN-2 (Figure 2A-B, Table S2). DDD107498 had the least effect on parasite growth during the treatment window, with mean growth inhibition of approximately 10%, 18%, and 24% in 1.8x-, 5x-, and 10x EC_50_-treated EEFs respectively (Table S2).

### Neither the speed and completeness of translation recovery, nor parasite growth after washout, is determined by the efficacy of translation inhibition

Our central hypothesis was that compounds capable of causing the most complete shutdown of parasite translation would result in stronger and more long-lasting effects on *P. berghei* liver stages after compound washout. To our surprise, this was clearly not true for translation recovery. The populations of parasites treated with anisomycin and LysRS-IN-2 recovered the fastest. 4 h after compound washout, these treatment groups were translating at levels comparable to the DMSO controls and had exceeded the magnitude of control translation by 44 hpi (Figure 3A). EEFs treated with bruceantin and DDD107498, the most and least effective translation inhibitors tested, showed nearly identical levels of translational output 4 h after compound washout (Figure 3A and Table S2). However, 16 h after washout, bruceantin-treated EEFs had completely recovered and even exceeded the control translational intensity, while those treated with DDD107498 showed little, if any, further recovery (Figure 3A). Parasites treated with 1.8x EC_50_ DDD107498 recovered substantially in 4 h, to approximately 67% of control translational intensity, with a near complete recovery to control levels by 44 hpi (Figure 3A and Table S2). MMV019266 treatment also had lasting effects on translation, with little recovery observed 4 h after washout, but on a population level, translation increased to ∼50% of DMSO controls after 16 h (Figure 3A). Looking at the individual data points for MMV019266 treated individual parasites at 44 hpi, recovery is quite heterogeneous, with some parasites translating at control levels while others remain completely inhibited (Figure 3A). Translation recovery in the HepG2 cells after anisomycin and bruceantin treatment largely paralleled the LS translation recovery observed (Figure S3), though bruceantin-treated cells did show greater recovery after 4 h, as would be expected given that bruceantin is somewhat *Plasmodium*-selective.

**Figure 3.**
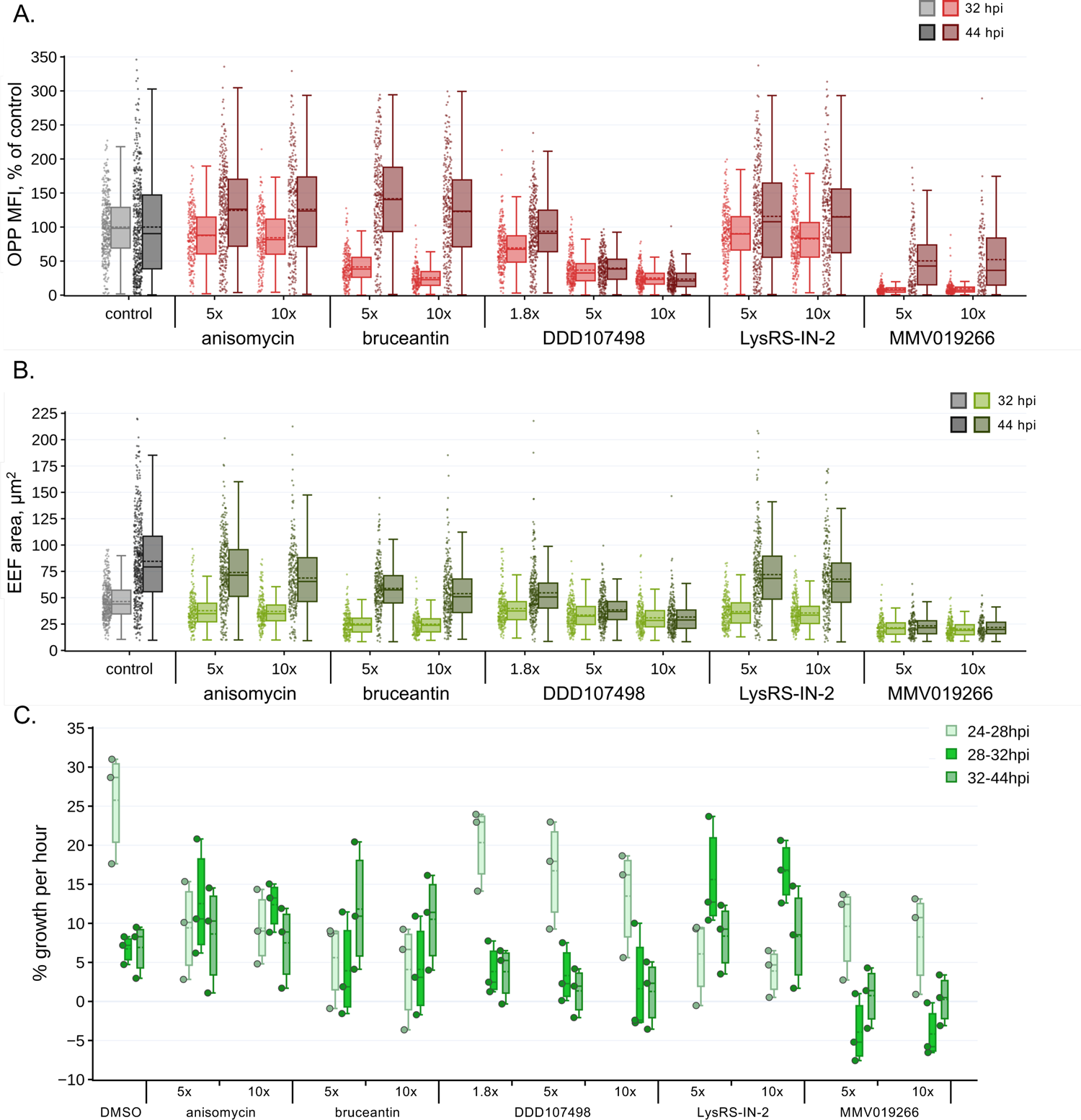
Post-washout translation recovery and growth of *P. berghei* liver stages. A) Parasite translation (OPP MFI, % of control), and B) parasite area (µm^2^) quantified 4 h (32 hpi) and 16 h (44 hpi) after compound washout, as schematized in Fig. 1C; individual points represent single EEFs. The full dataset for (A-B) can be explored via interactive dashboards in in our KNIME hub workflow: https://hub.knime.com/-/spaces/-/~EcnvMwYtqylu2reV/current-state/. C) Modeled % growth per hour during each time window, see Methods for details. Each point represents an experimental mean. Dotted lines in the boxplots represent the mean of n=3 independent experiments.

We next examined parasite growth after compound washout, in terms of parasite size at both recovery timepoints (Figure 3B); from this data, we also estimated % growth per hour for each condition tested, using the assumption that growth rates were linear and constant throughout each time interval for which we had measurements. In control parasite populations, the mean percentage of growth per hour from 24-28 hpi was inexpertly more than 4 times greater than during the 28-32 and 32-44 hpi intervals for all independent experiments, even though growth was variable from one experiment to another (Figure 3C). LysRS-IN-2 treated parasites had the fastest growth rates observed during the 28-32 hpi window, more than double that of the controls (Figure 3C), with anisomycin- and LysRS-IN-2 treated parasites reaching near equivalent sizes at 32 hpi (Figure 3B-C). Parasites from these two treatment groups continued to grow at a rate greater than or equal to that of the controls from 32-44 hpi (Figure 3B-C). This quick and profound growth is well-matched to the translation recovery dynamics observed for these compounds (Figure S4). In contrast, the bruceantin-treated parasites were growth impaired at 32 h, showing only a small increase in mean parasite size (Figure 3B). Just as with translation, bruceantin-treated parasites showed a large increase in size at 44 hpi and the percent of growth per hour during the 32-44 hpi window was the largest of any treatment group for both 5x- and 10x EC_50_ (Figure 3B-C, S4). Despite the robust growth of LysRS-IN-2, anisomycin, and bruceantin treated liver stages at both 5x and 10x EC_50_, parasites from all these groups remained smaller, on average, than controls at 44 hpi (Figure 3B and S4).

DDD107498-treated parasites were strikingly growth impaired after compound washout. EEFs treated with 5x EC_50_ grew slightly at both 32 and 44 hpi, equivalent to about half the growth rate seen in controls from 28-32 hpi, falling to <20% in the 32-44 hpi period (Figure 3B-C), while the size of EEFs treated with 10x EC_50_ DDD107498 changed very little from 28-44 hpi reflecting profound growth inhibition (Figure 3B-C). Even in the 1.8x-EC_50_ treatment, parasites grew slower than the controls, despite the translational recovery seen at both 32 and 44 hpi in the 1.8x EC_50_ DDD107498 treated parasites (Figure 3, S4). MMV019266-treated parasites showed no growth at all after compound washout (Figure 3B-C). Growth clearly lags behind or is uncoupled from translation for MMV019266-treated EEFs, as the large jump in translational intensity seen in 44 hpi parasites is not at all paralleled (Figure 3, S4). Our results show that growth does require resumption of translation, as would be expected. Taken together, these results demonstrate that translation inhibition efficacy does not determine either the speed or completeness of LS translation recovery timing nor the extent of *P. berghei* LS parasite growth after washout.

### A brief period of translation inhibition in early schizogony causes both developmental delay and developmental failure in *P. berghei* liver stages

The ultimate readout of LS developmental success in the *P. berghei-*HepG2 infection model is parasite maturation, culminating in the formation of hepatic merozoites and their release in from the monolayer inside detached cells. Across all treatment groups, late liver stage development is deleteriously impacted by a brief period of translation inhibition during early schizogony. In the normal course of LS development, the onset of hepatic merozoite surface protein 1 (MSP1) expression in the parasite plasma membrane begins in the late liver stage before cytomere formation and remains through hepatic merozoite formation (see e.g. (*32*)). Apical merozoite antigen 1 (AMA1) expression begin only as merozoites are being formed (Figure S5), and the release of hepatic merozoite-filled detached cells, the merosome-equivalent in the *in vitro* infection model, into the medium occurs last (*30, 33*). When parasite populations are analyzed at 72 hpi, there are notable impairments at each of these steps in parasites subjected to translation inhibition during early schizogony, but the level of impairment is compound and concentration dependent (Figure 4A). Late liver stage development was most similar in anisomycin and LysRS-IN-2 treatments (Figure 4A), though monolayers treated with anisomycin contained fewer hepatic merozoites (60-63% of control) compared to LysRS-IN-2 treatments (61-69% of control) (Figure 4A & Table S2). Monolayers treated with bruceantin showed less successful parasite maturation compared to those treated with anisomycin or LysRS-IN-2, particularly at the 10x EC_50_ concentration (Figure 4A). In the 5x- and 10x EC_50_ treatments with MMV019266 and DDD107498, very few parasites showed evidence of late liver stage maturation (Figure 4A). The average number of parasites in the MMV019266-treated monolayers was the lowest observed, suggesting that some of the treated parasites may be killed or cleared by the HepG2 cells. In the 1.8x EC_50_ DDD107498 treatments, MSP1-expressing EEFs were reduced to 50% of the controls, and hepatic merozoites abundance was only 17% relative to the controls (Figure 4A). As a whole, these results were again not compatible with translation inhibition efficacy determining downstream antiplasmodial effects.

**Figure 4.**
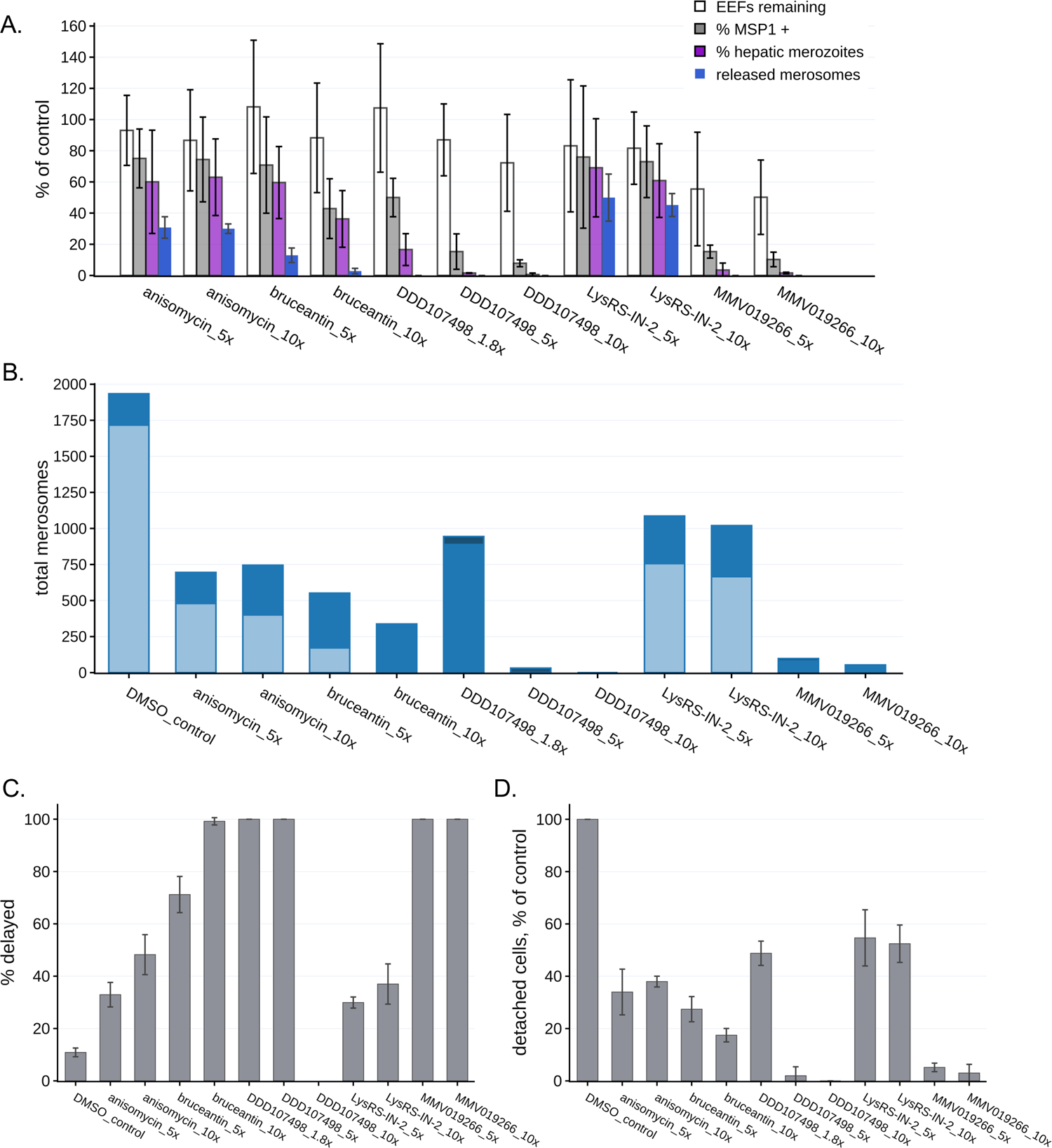
Translation inhibitor effects on late liver stage development. A) Parasite maturation at 72 hpi, normalized to DMSO control set at 100%, is quantified based on EEFs remaining in the monolayer, the % of monolayer EEFs expressing MSP1, the % of monolayer EEFs containing hepatic merozoites (expressing both MSP1 and AMA1, see Fig. S5), and merosomes released. Each treatment well contained a coverslip, which was used for monolayer maturation analysis and the medium contained detached merosomes which were collected and counted. Bars report the mean of n=3 independent experiments, error bars = standard deviation. B) Detached merosomes were collected and counted in experiment-matched wells at 72 (n=3 expts), 96 (n=3 expts), and 120 hpi (n=expts), with stacked bar charts showing the total merosomes per timepoint. C) The percent of total detached cells/ merosomes released after 72 hpi (% delayed) are reported as means with error bars showing standard deviation. D) Total detached cell/ merosome release normalized to the DMSO controls, with error bars showing standard deviation. For all panels, treatments are labeled by compound and multiple of EC50.

The harmful effects of a brief period of translation inhibition in early schizogony are even more apparent in 72 hpi merosome counts, with no merosomes collected from any of the DDD107498 or MMV019266 treated wells, and very few in the 10x EC_50_ bruceantin wells (Figure 4A). We anticipated that a 4 h period of translation inhibition would cause a small developmental delay, thus our matched assessment of monolayer and detached cells/merosomes at 72 hpi, but we also wanted to be able to separate more profound cytostatic effects from those that likely indicate developmental failure and compound cytotoxicity. Thus, we set up additional samples for repeated merosome harvesting from a single well, with collections made at 72-, 96- and 120 hpi (see Table S3 for merosome counts per experiment).

Control parasites that completed hepatic merozoite formation were largely released from the monolayer by 72 hpi, with no control merosome formation during the 96-120 hpi period (Figure 4B). Looking at total merosome counts over 3 days, every condition except 10x EC_50_ DDD107498 ultimately produced merosomes, with both LysRS-IN-2 treated samples producing the most (Figure 4B). Strikingly, the 1.8x EC_50_ DDD107498 treated-parasites were nearly equivalent to 5x- and 10x EC_50_ LysRS-IN-2 in overall merosome production, but the merosomes were only found in the 96 and 120 hpi collections (Figure 4B). Separating the data into cytostatic (% of delayed merosomes released after 72 hpi, Figure 4C) and potentially cytotoxic effects (% of total control merosomes, Figure 4D), we find that all treatments which led to any merosome production caused both cytostatic and cytotoxic effects on parasite populations. A clear difference can be seen between 5x- and 10x EC_50_ anisomycin treatments, with the later causing more cytostatic effects (Figure 4C), even though the eventual merosome production between the two was equivalent (Figure 4D). Despite inducing essentially total translation inhibition at both 5x- and 10x EC_50_ (Figure 2A), bruceantin was again the compound with the most variability between the 5x- and 10x EC_50_ treatments in terms of antiplasmodial effects, as cytostatic effects were lower and overall developmental success higher at the 5x EC_50_ concentration (Figure 4B-D). Brief treatment during early LS schizogony with MMV0192667 at both 5x- and 10xEC_50_ had extremely strong antiplasmodial effects, with overall developmental success of ∼5% and 3% respectively (Figure 4B, D), and though all merosomes were delayed, they formed in all 3 experiments (Table S3). Brief DDD107498 treatment at 5x- and 10x EC_50_ also appeared to overwhelmingly kill EEFs, with merosomes produced only after 96 hpi in a single experiment, and only in the 5x EC_50_ treatment (Fig 4B, D and Table S3).

### Correlations between translation inhibition efficacy and later *P. berghei* liver stage antiplasmodial effects are compound specific

Even with our limited compound set, it is clear that translation inhibition efficacy does not determine the speed or completeness of parasite recovery after a brief treatment with translation inhibitors. After compound washout, parasite translation, growth, and development appeared quite variable between compounds. To determine how well-correlated individual metrics were, both overall and for individual compounds, we performed correlation analysis of several key metrics describing the parasite response to acute translation inhibition. First, we examined correlations using data from all 5 compounds at both 5x- and 10x EC_50_. The only strong correlation seen across data from all compound treatments was a negative correlation between growth recovery from 28-32 hpi and growth recovery from 28-44 hpi (Figure 5A, see Table S3 for statistics). Analyzing the compounds individually, strong correlations that are compound specific are seen, with clear differences even for anisomycin and LysRS-IN-2 treatments, which were the two most similar compounds analyzed. For instance, in the LysRS-IN-2 treatments, there are strong positive correlations between translation recovery from 28-44 hpi and the MSP1 expression (correlation= 0.82, p= 0.001) and hepatic merozoites (correlation= 0.73, p= 0.007) (Table S3), while in the anisomycin treatments they are not correlated (translation recovery VS MSP1, correlation= 0.03; translation recovery VS hepatic merozoites, correlation= 0.10) (Table S3). Additionally, there is a weak, yet significant correlation between translation inhibition efficacy and growth inhibition at 28 hpi (correlation= 0.65, p= 0.02) in LysRS-IN-2 treatments, but not anisomycin (correlation= −0.07). This difference may be supported by the observation that on average, parasites grew more during the anisomycin treatments compared to the LysRS-IN-2 treatments (Figure 3C and Table S3), even though anisomycin was a more efficacious translation inhibitor (Figure 3A).

**Figure 5.**
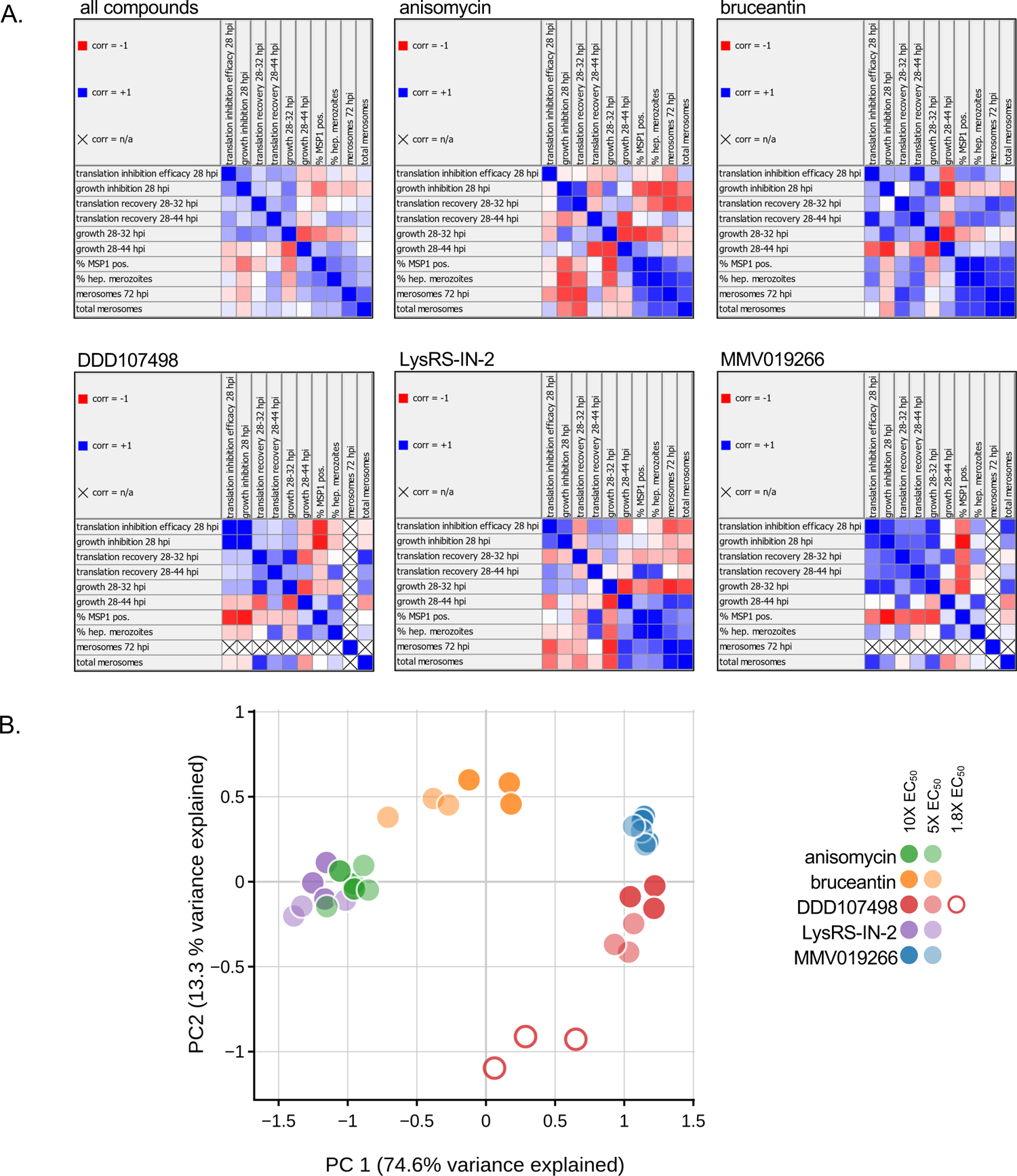
Multivariate analysis of translation inhibitor efficacy, recovery, and antiplasmodial effects on *P. berghei* liver stage parasites. A) Linear correlation matrices comparing compound efficacy, parasite recovery, and development (Pearson’s correlation, 2-sided; correlation values and p-values reported in table S2). Metrics used in correlation analysis were experimental means reported in table S2, or calculated from them, with 5x- and 10x EC50 concentrations combined (see Methods for details). Positions are marked X when one of the two metrics being compared has a value of 0. B) Principal components analysis using the same metrics analyzed in A), but with data separated by both concentration and individual experiments. The first two principal components explain a total of 87.9% of the variance.

When a principal components analysis is applied to the same set of features, but with data separated by both concentration and experiment, most compounds are clearly separated from one another, while anisomycin and LysRS-IN-2 are partially overlapping (Figure 5B). The 5x- and 10x EC_50_ treatments of MMV019266 were essentially indistinguishable, and also had the greatest similarity between experiments of the compounds tested (Figure 5B). Overlap is also seen for the two anisomycin and LysRS-IN-2 concentrations, but not for 5x- and 10x EC_50_ bruceantin or DDD107498 treatments (Figure 5B). Unsurprisingly, the 1.8x EC_50_ DDD107498 datapoints are distant from the other DDD107498 concentrations (Figure 5B).

### DDD107498 exerts antiplasmodial effects on translationally arrested parasites

Though DDD107498 at both 5x- and 10x EC_50_ concentrations was the least efficacious translation inhibitor tested, it had the strongest antiplasmodial effects. Strikingly, the 1.8x EC_50_ concentration (20 nM), caused only approximately 60% reduction in parasite translation output and the least growth inhibition during the treatment period, yet had stronger effects on translation recovery and growth after washout (Figure 3 and S4) and in LS developmental success (Figure 4), than 5x EC_50_ concentrations of anisomycin and Lys-RS-IN2. Given that *in vitro* antiplasmodial potency of DDD107498 is consistently between 1-2 nM in (Table S1 and (*5*)), we tested whether DDD107498 achieves a more profound LS translation inhibition after prolonged treatment. Treatment with 20 nM DDD107498 for 24 h, from 24-48 hpi, did not lead to increased LS translation inhibition (Figure S6). Taken together, this data led us to wonder whether DDD107498 can exert antiplasmodial effects independent of its ability to inhibit translation. To test this, we designed an experiment to determine if DDD107498 can exert antiplasmodial effects on translationally arrested *P. berghei* LS parasites. A supra-maximal concentration of anisomycin (10 μM, equivalent to 38x the translation inhibition EC_50_) was added at to *P. berghei* LS at 24 hpi to shut down translation, followed by addition of 20 nM DDD107498 at 25 hpi, thorough compound washout at 28 hpi, and analysis 20 h later at 48 hpi (Figure 6A). We have previously demonstrated that LS parasites recover from this anisomycin treatment regimen in terms of both translation and growth, and that 10 μM anisomycin treatment for 30 minutes is sufficient to cause essentially complete inhibition of *P. berghei* LS translation (*17*). Strikingly, anisomycin-arrested parasites subsequently treated with DDD107498 were smaller on average and had lower mean translational output then either anisomycin or DDD107498 single treatment controls (Figure 6B-C). We next used the same treatment regimen, but read out antiplasmodial efficacy with merosome collections at 72-, 96-, and 120 hpi (Figure 6D). Anisomycin-arrested EEFs subsequently treated with DDD107498 had a marked decrease in merosome formation with respect to the DMSO, anisomycin, and DDD107498 controls (Figure 6E-G, Figure S7, Table S4). A 3 h DDD107498 treatment gave a highly similar merosome production profile (Figure 6E) compared to the 4 h treatment (Figure 4B). DDD107498-treated anisomycin-arrested parasites showed a near identical cytostatic effect as those treated with DDD107498 alone, and completely distinct from those treated only with anisomycin (Figure 6F). Essentially all the difference between DDD107498 treatment of translating vs. anisomycin-arrested parasites was reflected in total merosome production up to 120 hpi (Figure 6G), reflecting a likely increase in parasite death. Given these unambiguous results, we repeated these experiments using 5x translation inhibition EC_50_ concentrations of the three other translation inhibitors studied (Table S1) to shut down parasite protein synthesis before addition of DDD107498, while maximizing the difference in antiplasmodial profile of each compound alone vs. that of DDD107498. In each case, addition of DDD107498 to the translationally arrested parasites led to 100% delayed merosome release, with LysRS-IN-2- and bruceantin-treated parasites shifted to the phenotype of DDD107498-treated parasites (Figure 6H-I, Figure S7, Table S4). Addition of DDD107498 to translationally arrested parasites exerted a remarkably clear and stable increase in likely parasite killing, with at least 50% reduction in total merosomes formed, regardless of whether anisomycin, bruceantin, LysRS-IN-2, or MMV019266 was used to inhibit parasite protein synthesis (Figure 6H, J).

**Figure 6.**
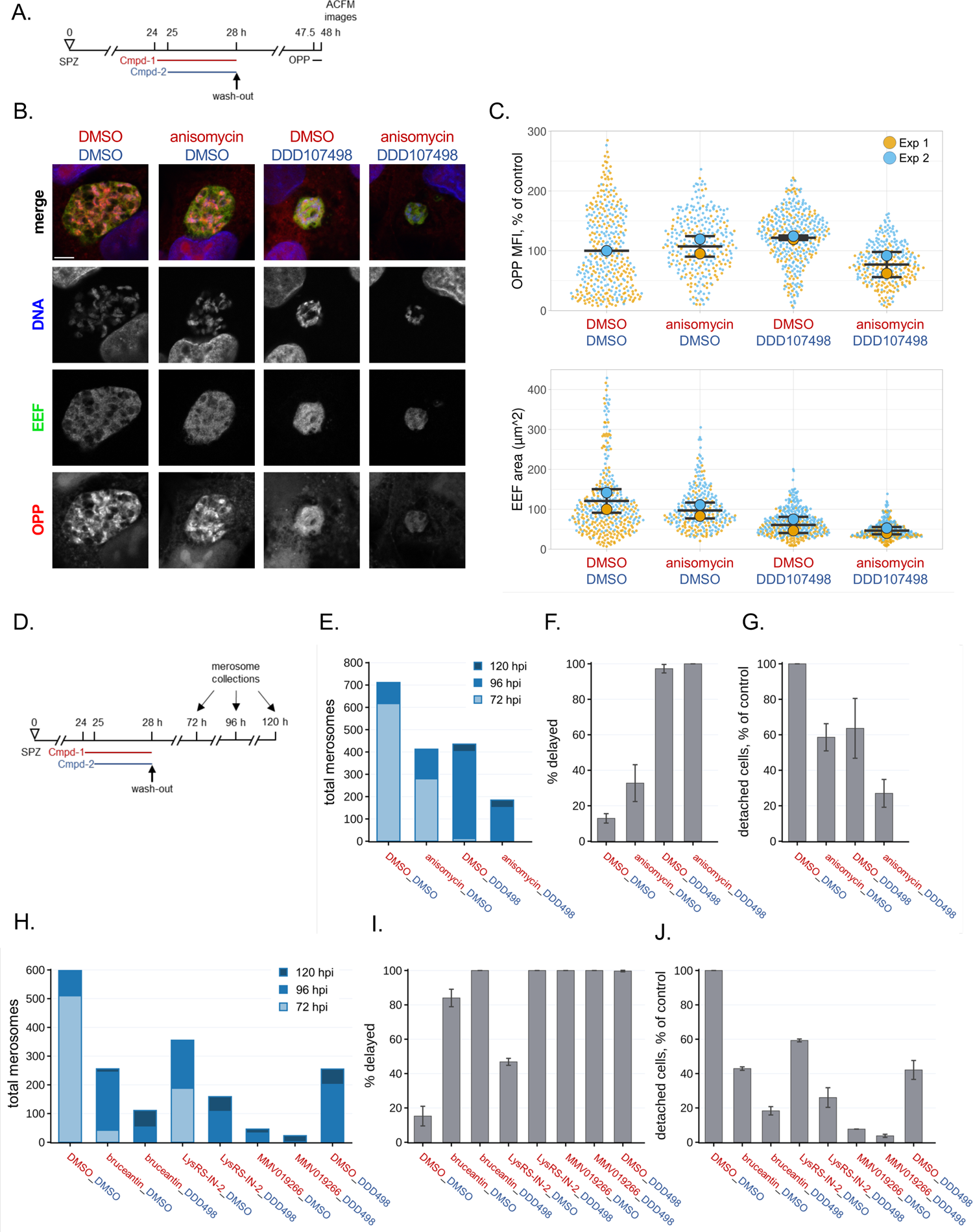
Antiplasmodial effects of DDD107498 on translationally-arrested *P. berghei* LS parasites. Legend on following page. Figure 6. Antiplasmodial effects of DDD107498 on translationally-arrested *P. berghei* LS parasites. A) Experimental schematic for panels B-C quantifying translation recovery and growth at 48 hpi after DDD107498 treatment of anisomycin-arrested parasites in early schizogony. B) Representative single confocal images of translation in *P. berghei* LSs at 48 hpi. Merged images are pseudocolored as indicated with parasite (EEF) immunolabeled with α-HSP70, OPP-A555 labeling the nascent proteome, and DNA stained with Hoechst. Scale bar = 5 μm. C) Single parasite translation and size quantified at 48 hpi. Single points show individual EEFs color coded by independent experiment, with experimental means represented by large circles (n=2), and bars represent mean and standard deviation of the experiments. D) Experimental schematic for quantifying merosome release in experiment-matched wells at 72, 96, and 120 hpi, after DDD107498 treatment of translationally arrested early LS schizonts and subsequent washout. Stacked bar charts (E,H) report total merosomes collected at per timepoint from all experiments. F,I) The percent of total detached cells/ merosomes released after 72 hpi (% delayed) are reported as means with error bars showing standard deviation. G,J) Total detached cell/ merosome release normalized to the DMSO controls, with error bars showing standard deviation. Data shown from n=3 (E-G) or n=2 (H-J) independent experiments; DDD498 = DDD107498 in the axis labels for (E-J).

## Discussion

Translation inhibitors, acting via diverse molecular targets to abrogate cytoplasmic *Plasmodium* protein synthesis, are attractive antimalarial compounds for drug development, as they should have multistage antimalarial activity, and thus be capable of treating symptomatic infection in the blood and providing chemoprotection against the asymptomatic liver infection. Here, we examined the relationship between translation inhibition efficacy and antiplasmodial efficacy in the *P. berghei* LS, treating parasites with five mechanistically distinct translation inhibitors at equivalent effective concentrations for 4 h in early schizogony, then systematically assessed the effects this treatment had on the parasites over 4 days. These experiments generated many intriguing results worthy of follow-up study and the firm conclusion that the extent of translation inhibition a compound induces does not determine the strength of its downstream antiplasmodial effects. The most dramatic demonstration of this is that the least efficacious translation inhibitor tested, DDD107498, had the strongest antiplasmodial effects at both 5x- and 10x translation inhibition EC_50_ concentrations. Even excluding DDD107498, which has the greatest disconnect between LS biomass inhibition EC_50_ and translation inhibition EC_50_ of the three *Plasmodium*-specific inhibitors tested, data from anisomycin, bruceantin, LysRS-IN-2, and MMV019266 also strongly demonstrate that antiplasmodial efficacy is not determined by translation inhibition efficacy. Bruceantin shows no difference in translation inhibition efficacy at 5x-vs 10x EC_50_ concentrations, but clear differences in antiplasmodial outcomes downstream of compound washout are seen. MMV019266 has much greater *P. berghei* LS antiplasmodial efficacy than anisomycin at both 5x- and 10x EC_50_, despite being a clearly less efficacious translation inhibitor. At 10x EC_50_, LysRS-IN-2 is a more efficacious translation inhibitor than MMV019266, but a much less efficacious antiplasmodial inhibitor. Clearly, translation inhibition efficacy alone is not a suitable metric to predict the strength of antiplasmodial efficacy of candidate compounds. Correlation and principal components analyses suggested that each compound tested has a specific antiplasmodial effect profile downstream of the translation inhibition it induced, with no metric strongly correlated to merosome production across all the compounds. Mechanism also does not appear likely to be informative, though our test set was quite small. LysRS-IN-2 and anisomycin were by far the two most similar compounds across all metrics we analyzed, despite having no mechanistic similarity whatsoever upstream of their effects on protein synthesis. Anisomycin inhibits translation elongation (*34*), and structural data in other eukaryotes indicates that it binds to the A-site of the 60s ribosome subunit (*22*), while LysRS-IN-2 targets the *P. falciparum* lysyl tRNA synthetase (Pf cKRS), which catalyzes aminoacylation of lysine onto its cognate tRNA for incorporation into nascent polypeptides, by binding within the enzyme’s ATP binding pocket (*28*). Taken together, our data suggest that the no single metric can provide an adequate correlate of LS antiplasmodial effects of translation inhibitors; instead, quantification of total merosome production up to 120 hpi is needed. Quantifying merosome production unfortunately relies on manual counting, making it both time consuming and error prone. Reproducible and automated assays that allow for merosome, or better yet, individual hepatic merozoite quantification will be needed.

Amongst our surprising findings in this work was the strength of antiplasmodial effects exerted by a 4 h compound treatment during early LS schizogony. We chose the 4 h treatment based on previous findings (*17*) that: 1) these compound all achieve profound translation inhibition within 30 minutes, suggesting that a 4 h treatment ensures several hours of maximal inhibition, and 2) high concentration anisomycin treatment for 4 h was reversible, in terms of both translation and parasite growth, after washout. We did not anticipate that the weakest effect we would observe on parasite developmental success after compound washout would be ∼50% reduction in release of hepatic merozoite-filled merosomes/detached cells. This indicates substantial heterogeneity in individual parasites’ abilities to withstand drug pressure, and a deeper understanding of which individuals are killed by the most reversible compounds may yield valuable insights.

The speed with which antimalarial drugs can kill parasites is a key consideration for therapeutic use, since speed in reducing parasite load can directly influence clinical outcomes. In the *P. falciparum* ABS, speed-of-kill is determined using parasite reduction ratio (PRR) assays (*27, 35*), in which mixed stage parasite cultures are subjected to compound treatment for 120 h, with aliquots drawn every 24 h that are thoroughly washed, and then put back into compound-free culture with limiting dilution. Cultures are then monitored for several weeks to detect parasite recrudescence and calculate the number of viable parasites present. When compounds are tested at equivalent effective concentrations, the PRR assay reveals compound specific differences that are masked in 48- or 72 h growth assays, where compounds often appear equally effective (*36*). This is also true for standard liver stage assays that quantify EEF biomass (*37, 38*), typically after 48 h of continuous treatment starting about the time of sporozoite addition, where any compound active against the EEF at the earliest stage of development will present a similarly dramatic reduction in biomass. Parasites treated with saturating concentrations of DDD107498, LysRS-IN-2, or MMV019266 all have such low biomass values in our 48 h live luciferase assay (ranging from 0.7%-1.5% of the DMSO control) that they fall within the background noise from an empty well. As the LS does not cycle, a PRR-like assay is challenging to implement. Viability must be assessed in the same parasites treated, not their progeny, in the absence of an integrated liver-to-blood infection model, and the equivalent of an ABS mixed culture is not easily realized. Given the remarkable phenotypic separation we observed between EEFs briefly treated with equivalent effective concentrations of different translation inhibitors during early schizogony, it will be worth investigating if this testing modality can provide useful insight across compound classes into liver stage speed-of-kill (or perhaps speed-of-effect), with the clear caveat that we cannot be certain that parasites which fail to release merosomes by 120 hpi are truly non-viable. As stage specificity will exist for LS-active compounds as well, it might be worthwhile to run parallel assays targeting (e.g.) 4-8 hpi and 48-52 hpi, so that early and late LS activity are also captured in the *P. berghei-*HepG2 infection model. Fo*r* the compounds inhibiting translation, at least, the early schizogony treatment workflow is quite informative. It is notable that the two aaRS inhibitors, LysRS-IN-2 and MMV019266 display the same profound difference in speed-of-effect/kill as observed in PRR assays where 10x growth inhibition EC_50_s were tested. LysRS-IN-2 is slower to kill mixed *P. falciparum* ABS cultures than even atovaquone, the slowest-acting reference compound, while MMV019266 and similar thienopyrimidines kill as quickly as artemisinin, the fastest-acting reference compound (*26, 28*). LysRS-IN-2 was the most reversible compound we tested, and yet still able to reduce merosome release by ∼50% at both 5x- and 10x translation inhibition EC_50_, while MMV019266 exerted much greater antiplasmodial effects, reducing developmental success to ∼ 5% at both 5x- and 10x EC_50_, with substantial developmental delay. DDD107498 tested at 5x- and 10x translation inhibition EC_50_ was the most efficacious antiplasmodial compound of the 5 translation inhibitors tested; with not a single merosome recorded at the 10x EC50 concentration in any experiment. This is not in keeping with its *P. falciparum* PRR profile, which is slow-killing like atovaquone, however (*5*).

DDD107498 presents a rather puzzling case as a translation inhibitor. We detected no LS translation inhibition activity from 1 nM DDD107498 in our earlier study, and only ∼40% inhibition at a 10 nM concentration (*17*), roughly the lower end of a 10x LS biomass EC_50_ estimate. This disconnect is not liver stage or *P. berghei-*specific. Using a *P. falciparum* ABS lysate-based approach, the translation inhibition IC_50_ for DDD107498 was determined to be 60.5 nM in the PfIVT assay (*19*), while testing at 10x the antiplasmodial EC_50_ led to ∼45% or ∼70% translation inhibition using ^35^S-labelled amino acid incorporation assays in *P. falciparum* ABS populations (*5, 19*). Because of the nearly 10-fold increase in translation inhibition EC_50_ vs. LS biomass inhibition EC_50_ for DDD107498, and its place amongst promising new antimalarial compounds progressing in clinical trials (*6, 7*), we also ran DDD107498 at 20 nM – roughly, an upper estimate of 10x EC_50_ for LS biomass inhibition, as this concentration supports more direct comparison with other data on the compound’s antiplasmodial efficacy. The 20 nM concentration (1.8x LS translation inhibition EC_50_) induced ∼60-65% translation inhibition, in keeping with the shallow slope of the fit concentration-response curve. LS parasite growth inhibition was maximal after compound washout, and translation recovered slowly, in comparison with compounds like anisomycin or LysRS-IN-2 which caused far more profound translation inhibition. 1.8x EC_50_ DDD107498 treatment induced stronger antiplasmodial effects than LysRS-IN-2 at 5x- and 10x EC_50_, and all DDD107498-treated parasites that formed merosomes displayed a marked cytostatic effect. 1.8x DDD107498-induced phenotypes were so different from those of 5x- and 10x EC_50_, that it appears to be essentially a distinct compound in the PCA analysis. Given these results, we designed an experiment to ask if 20 nM DDD107498 could exert antiplasmodial effects on EEFs translationally arrested by a super-saturating concentration of anisomycin for an hour before DDD107498 addition. Not only could addition of DDD107498 to these translationally arrested parasites (∼99% translation inhibition expected, per the 10x EC_50_ data) induce the same cytostatic effects observed when translating parasites were treated, the staggered combination treatment reduced merosome formation by roughly 50% compared to anisomycin treatment alone. Anisomycin was chosen for this analysis because it maximized the difference between single treatment-induced merosome effects, but on the strength of this result, we found that DDD induced a similar ∼50% reduction in developmental success in parasites translationally arrested by the other three compounds in our study. We note that these experiments were not designed to test for synergy between compounds, and are insufficient to do so, but we interpret the data as more suggestive of additive effects. It will be worthwhile to look at this more carefully, as the combination of multiple active compounds into a single antimalarial medicine is critical for preventing blood stage resistance from developing (*11*), but may also be able to effect better chemoprotection against the LS. Unexpected synergy between bacterial translation inhibitors has been documented (*39*), and it seems worthwhile to test for such synergy between antiplasmodial translation inhibitors in the LS, though the merosome release readout is problematic to scale.

Baquero and Levin (*40*) provide a useful framework emphasizing the distinction between proximate vs. ultimate causes of antibacterial action by antibiotics, and it is worthwhile to consider antiplasmodial activity, in particular, that of DDD107498, in this light. Is the proximate cause of DDD107498 antiplasmodial effects against LS parasites mediated by a target other than eEF2? This seems quite unlikely, given the centrality of eEF2 to protein synthesis and the overwhelming similarity of DDD107498 potency across species and stages (*5*), but direct examination of LS parasites with DDD107498-resistant eEF2 alleles will ultimately be needed. The genetic evidence linking eEF2 mutations to drug resistance is extremely strong, and no other parasite genes have been additionally implicated to date (*5, 6, 20, 21*); further, a mutation that shifts *P. falciparum* ABS antiplasmodial potency also shifts *P. falciparum* ABS translation inhibition potency (*5*). It is notable that eEF2 has roles beyond catalysis of the translocation step of elongation, which are also potential proximate causes of DDD107498 activity. eEF2 is involved in the fidelity of ribosomal translocation via its diphthamide modification (*41, 42*), and catalyzes reverse translocation (*43*). In the presence of a compound able to stabilize eEF2 on the ribosome, as has been demonstrated for sordarin (*44*), the likelihood of reverse movement is increased, which has been suggested to contribute to sordarin’s activity (*43*). Though we detect substantial translation in the presence of 20 nm DDD107498, the makeup of the nascent proteome under these conditions is unknown. If translation fidelity or processivity are affected, the polypeptides synthesized might be nonfunctional and sufficiently aberrant to induce cellular stress, a possible ultimate cause of antiplasmodial activity that is largely consistent with our observations. As we show that prolonged incubation with 20 nM DDD107498 does not lead to increased translation inhibition in *P. berghei* EEFs, the hypothesis that translation inhibition becomes stronger with prolonged treatment can be rejected. However, the argument against DDD107498 activity being ultimately due to a toxic nascent proteome is hard to reconcile with the fact that DDD107498 exerted antiplasmodial activity against parasites that were translationally arrested during the entirety of their exposure to the compound. This could possibly be explained if DDD107498 was retained in the parasite after compound washout, perhaps in a stable complex with binding partners. DDD107498 could also exert LS antiplasmodial effects on processes in addition to protein synthesis, via hypothetical non-canonical roles of eEF2 (as described for other elongation factors (*45, 46*)) or via additional targets.

Antibacterial translation inhibitors have been extensively studied (*47, 48*), yet far less is known about their ultimate killing mechanisms. It has been suggested that bactericidal antibiotics, including those that target the ribosome, ultimately kill bacteria by stimulating production of hydroxyl radicals, which bacteriostatic antibiotics, regardless of mechanism, do not (*49*), though this has been controversial (*50*). Antibiotics targeting protein synthesis fall into both bactericidal and bacteriostatic categories, sometimes even within the same class of molecules, such as the macrolides, for which evidence exists that cidality is linked to slow dissociation rates from the ribosome (*51*). For aminoglycosides, clinical mainstays which inhibit protein synthesis by binding with high affinity to the ribosomal A site (*52*), several hypotheses for ultimate cause of bacterial killing have been proposed, including ribosome independent but not alternative direct effects on the bacterial outer membrane (reviewed in (*40*)). Comparable compound-specific systemic effects leading to parasite killing could be at play for antimalarial translation inhibitors, as far less is known about their ultimate parasite killing mechanisms. Our data as whole suggest these ultimate killing mechanisms are worthy of further study, as the hypothesis that *P. berghei* LS antiplasmodial killing by translation inhibitors is determined by translation inhibition efficacy *per se* should be rejected.

The rate of cellular growth is influenced by cell-extrinsic factors like nutrient availability, along with many intrinsic factors, like protein concentration, ribosomal capacity, the rate of protein synthesis and cell size (*53–56*). In organisms where replication culminates in binary fission, including several bacterial, fungal, and metazoan species, cellular growth rates have been observed to be exponential (*53, 55, 57*), where absolute growth is directly proportional to cell size, though this remains somewhat controversial (*58*). We used an assumption of constant, linear growth to estimate % growth per hour for *P. berghei* EEFs based on data collected at 24-, 28-, 32-, and 44 hpi, which led to the unexpected finding that % growth per hour was ∼4 times greater during the 24-28 hpi period, than for the other two. Maintaining an appropriate nuclear-to-cytoplasmic ratio is crucial for growth and cellular functions (*59, 60*); since both DNA content and cell size increase immensely during *Plasmodium* EEF growth, there should be no reason to assume that growth must slow as the parasite grows. While our data is sparse, it does suggest that growth is not constant, and that the *P. berghei* LS cellular state may be front-loaded to achieve rapid growth earlier in development. Despite the variability in growth rates and translational intensity seen between independent experiments, which likely represents biological variation in sporozoite fitness between infections, the growth rate variability between timepoints was robustly observed in each experiment. While we were not able to identify a technical explanation for the differences in growth rates, particularly since some compound-treated samples exhibited high growth rates after washout in both the 28-32 and 32-44 hpi windows in which control growth was slow, experiments purposely design to query LS parasite growth rates over time will be needed to validate our preliminary findings.

Due to the high levels of translation inhibition efficacy we observed at 28 hpi (>90% for all compounds except DDD107498) and evidence that most compounds reach similar levels of inhibition after only 30 minutes of treatment (*17*), it is notable that parasite growth inhibition was not more profound during the 4 hours in which the compound was present; none of the compounds reduced growth from 24-28 hpi by even 50%. This could suggest that a small amount of residual translation is enough to support some parasite growth, or there may be a temporal lag in parasite growth during this stage in development. Alternatively, parasite growth may be driven by a high steady-state protein concentration at the onset of treatment. In fission yeast, cellular growth appears to function as a mechanism controlling intracellular protein concentrations (*55*). If yeast cells were prevented from increasing in size while protein synthesis continued at a rate equivalent to log growth, thus artificially increasing the concentration of protein in the cells, the yeast entered a phase of hyper growth when released from the growth constraints. Interestingly, LysRS-IN-2 induced what could be interpreted as a hypergrowth phenotype immediately following compound washout; the % growth per hour of LysRS-IN-2-treated parasites during the 28-32 hpi window was nearly as high as the controls exhibit during the 24-28 hpi period. Bruceantin-treated parasites remained profoundly growth inhibited for the first 4 h after washout despite partial recovery of translation during the same time period but had the fastest growth rate during the 32-44 hpi window. This perhaps suggests that the cellular state at 24 hpi supporting rapid growth in control parasites is maintained in those that are profoundly translationally arrested at that time and can be harnessed for a period of quick growth after compound washout, once translation levels are sufficient. This raises the question of how much translation is necessary to support *P. berghei* LS growth. Parasites treated with 5x- or 10x (translation inhibition) EC_50_ DDD107498 show little growth between 32 and 44 hpi, despite having ∼37% or 24% of control translational output respectively, suggesting this amount is not sufficient. Parasites treated with 1.8x EC_50_ DDD107498 showed about 36% residual translation at 28 hpi, which recovered to ∼67% of control translation by 32 hpi and 94% of the control by 44 hpi, yet parasites continued to grow at a slower rate than the controls, with the largest decrease seen 32-44 hpi. Finally, MMV019266-treated EEFs, as a population, have reached close to 50% of control translation at 44 hpi, and many individuals are translating at normal levels, but this does result in any detectable growth in individuals or at the population level. Analysis after 44 hpi would be needed to understand if and/or when growth is observed in response to the resumption of translation for these parasites. As a whole, this data once again suggests that compound specificity may be at play, but more work is needed to gain a comprehensive understanding. A limitation of our work is that the effects downstream of translation inhibition may be specific to the parasite developmental stage, particularly as the control growth rate we observed was so much higher from 24-28 hpi than in subsequent periods. Further work will be needed to assess how generalizable these results are across different treatment windows during the *P. berghei* LS and those of the clinically relevant *Plasmodium* spp. Considering the magnitude of growth that occurs without any cell division until the hepatic merozoite progeny are formed in a mass cellularization event, the *Plasmodium* liver stage parasite will be a fascinating system for further studies of growth mechanisms and their integration with translational output.

## Materials and Methods

### HepG2 cell culture and infection by *P. berghei* sporozoites

Human hepatoma (HepG2) cells were cultured in complete Dulbecco’s Modified Eagle Medium (cDMEM) (Gibco 10313-021) supplemented with 10% (v/v) FBS, 1% (v/v) GlutaMAX (Gibco 35050-061), 1% (v/v) Penicillin-Streptomycin (Gibco 15140-122) and maintained at 37°C, 5% CO_2_. Sporozoites were isolated from *Plasmodium berghei-*infected *Anopheles stephensi* mosquitos (New York University Insectary and University of Georgia SporoCore); sporozoite isolation and infection of HepG2 cells were performed as previously described (*17*). Briefly, *P. berghei* sporozoites (ANKA 676m1cl1 from BEI Resources, MRA-868) with dual reporter expression construct (exoerythrocytic form 1a (EEF1a) promoter controlled firefly luciferase and GFP fusion protein (*61*)) were extracted from *Anopheles stephensi* mosquito salivary glands, counted and diluted into complete DMEM further supplemented with 1% (v/v) Penicillin-Streptomycin-Neomycin (Gibco 15640-055), 0.835µg/mL Amphotericin B (Gibco 15290-018), 500µg/mL kanamycin (Corning 30-006-CF), and 50µg/mL gentamycin (Gibco 15750-060) (iDMEM), before being exposed to HepG2 monolayers, centrifuged at 3000 rpm for 5 minutes and incubated, as described, for 2 hours.

After incubation, monolayers were washed with PBS and replenished with iDMEM for direct infections occurring in 24 well plates with or without glass coverslips. For experiments carried out in 96 well plates (Greiner 655098), infected cells were detached using TrypLE Express (Gibco 12605-028), washed, counted, and re-seeded at the desired density. All *P. berghei*-infected HepG2 were maintained at 37°C, 5% CO_2_. Live luciferase activity following 48 h treatment was performed as previously described (*25*).

### OPP labeling, compound treatments and washouts

20 µM O-propargyl puromycin (OPP) (Invitrogen C10459) was used to label cells for 30 mins at 37°C, before 15-minute fixation with paraformaldehyde (PFA) (Alfa Aesar 30525-89-4) diluted to 4% in PBS at RT. Anisomycin (EMD Millipore / Sigma, 176880), bruceantin (MedChem Express, HY-N0840), DDD107498 (Apex Bio, A8711), Lys-RS-IN2 (MedChem Express, Y-126130), and MMV019266 (Vitas-M Laboratory, STK845176) were solubilized in DMSO (Sigma-Aldrich D2650), aliquoted, and stored at −20°C. Acute pre-treatment assays were performed as previously described (*17*). Briefly, compounds were applied for 3.5 hours followed by 30-minute OPP labeling in the presence of compound treatment. For the competitive OPP (co-OPP) assay performed in Figure S1, LysRS-IN-2 and OPP were applied simultaneously for 30 minutes. Equimolar concentrations of DMSO (0.1% − 0.2% v/v) were maintained for all DMSO controls and treatments. For samples to be analyzed after 28 hpi, samples were subject to a stringent compound washout protocol, where the wash volume and total number of media exchanges was calculated to ensure a ≥ 4-log reduction in compound concentration per well. At the end of the 4 h treatment, fresh iDMEM was added to each well to triple the final volume, reducing 10x EC_50_ concentrations to 3.3x EC_50_. Next, all media was removed by pipette and replaced with fresh iDMEM, for a total volume of 1200µL per well in 24 wp, and 300 µL per well in 96 wp. This wash step was repeated for a total of 3 full washes. Based on the average retained iDMEM volume in each well type (80 µL in 24 wp, and 20 µL in 96 wp) the 10x EC_50_ treatment should be exposed to ≤ 0.001x EC_50_ after the last wash.

### Click Chemistry and Immunofluorescence

Click chemistry was performed using copper-(I) catalyzed cycloaddition of Alexafluor555 picolyl azide using Invitrogen Click-iT Plus AF555 (Invitrogen C10642), largely according to the manufacturer’s protocol. All click reactions were performed using a 1:4 Cu_2_SO_4_ to copper protectant ratio. *P. berghei* infected HepG2 cells were immunolabeled using anti-PbHSP70 (1:200, 2E6 mouse mAb) (*62*)to mark EEFs followed by donkey anti-mouse Alexafluor488 (Invitrogen A21202). MSP1 was immunolabeled with Anti-PBANKA_083100, affinity-purified rabbit polyclonal (peptide SSTEPASTGTPSSGC, produced by GenScript, 1:1000), followed by donkey anti-rabbit Alexafluor555 (Invitrogen A31572) and AMA1 was labeled using *P. falciparum* rat monoclonal 28G2 (*63*) (MRA-897, BEI Resources, 1:300), followed by donkey anti-rat Alexafluor647 (Invitrogen A48272). All secondary antibodies were used at a concentration of 1:500. DNA was stained with Hoechst 33342 (Thermo Scientific 62249) (1:1000). All labeling was carried out in 2% BSA in PBS.

### Image acquisition

ACFM imaging was performed using a Leica SP8 confocal microscope using an HC PL APO 63x/1.40 oil objective for glass coverslips (Figures 6B-C; S1A,B,D; and S5), or an HC PL APO 63x/1.40 water objective for 96-well µclear plates. Images in Figure S1D were acquired manually and were processed using Fiji (*64*). For Figure 4A, HCI images were acquired using a 10x air objective. All other images presented in manuscript, as wells as images used for quantitative analysis were acquired using automated confocal feedback microscopy (ACFM) as previously described (*17, 29*).

### Image segmentation, feature extraction, and data cleaning

Batch image segmentation and feature extraction were performed using CellProfiler (version 2.1.1 rev6c2d896) (*65*), as previously described (*17*). Briefly, for ACFM images, EEF objects were segmented using a global Otsu thresholding strategy of the PbHSP70 image. Within each image set, the EEF object was either shrunk or expanded by two pixels for subsequent image masking to ensure exclusion of HepG2-associated signal in masked OPP-A555 images used for intensity feature extraction. DNA objects within EEF objects were also identified by global thresholding and unified based on known parasite sizes. All HepG2 nuclei identified in an image were unified into a single object, and the OPP-A555 fluorescence intensity features of this unified object were used to quantify specific signal associated to HepG2 translation. Size, shape, and fluorescence intensity features of both EEFs and in-image HepG2 were exported from CellProfiler for downstream analysis using KNIME (*66*). To ensure the analysis included data from ACFM image sets containing one (and only one) true EEF in a HepG2 monolayer, as opposed to a piece of fluorescent debris or a dying cell not integrated in the monolayer, we computationally removed all data associated to image sets in which: more than one EEF object was identified, the EEF object identified did not contain a DNA signal, and no HepG2 nuclei were identified. Additionally, out of focus EEFs were identified and removed using object location filters in X and Y planes. Finally, metrics for EEF object area, EEF object circularity (compactness), and the identification of unified EEF DNA objects were used to identify cases of segmentation failures in which two parasites were segmented as a single EEF object. Our workflow for quality control of ACFM images can be accessed at https://hub.knime.com/-/spaces/-/~TZCrKvv3sbJwM_xP/current-state/.

### Merosomes/detached cells collection and counting

For merosome/detached cell collections in Figures 4 and 6, the 1 mL iDMEM from each infected well was collected and transferred to a 24-well plate, then 500 μL of iDMEM was gently added to each infected well, then collected and pooled with the initial 1 mL, and the merosomes/detached cells were allowed to settle for 15 minutes before being counted a light microscope equipped with an X-Y adjustable stage at 40x magnification. For collections in figure 4B and 6D-J, 1 mL of iDMEM was added back to each infected monolayer and incubated for an additional 24 hrs, with merosomes/detached cells recovery and counting was repeated at the 96- and 120 h timepoints.

### Data analysis, concentration response curve fitting and statistics

For concentration-response analysis, 4 parameter non-linear regression curve fitting was performed in GraphPad Prism (Version 7.0d) with the minimal response (top of the curve) set at 100, a constrained hill slope (−10< hill slope< 0). For analyses in which maximal effect was reached with ≥2 consecutive concentrations, the maximal effect was fit open; if no such plateau was achieved, the curve was fit with maximal effect constrained to 0. EC_50_ and 95% CI were determined for each compound from ≥ 3 independent experiments. One way ANOVA with Tukey’s multiple comparisons post hoc testing were performed in GraphPad Prism (Version 9.5.1) using the mean of 3 independent experiments. All other data and statistical analyses were performed in KNIME (Version 4.5.1). In Figure 3C we modeled % growth per hour, with the assumption that growth would be constant throughout the time period analyzed, by determining the percent growth (experiment mean EEF area) from one timepoint to another, divided by the number of hours separating the two measurements, e.g. (((28 hpi area – 24 hpi area)/24 hpi area)*100)/4) would yield % growth per hour between 24 and 28 hpi. Plots were constructed using KNIME, Plotly Chart Studio, and the SuperPlotsOfData online tool (https://huygens.science.uva.nl/SuperPlotsOfData/).

## Data Availability

In addition to the supplementary tables, the complete dataset used to generate Figure 1A-B can be downloaded and explored via interactive dashboards via our KNIME hub workflow at https://hub.knime.com/-/spaces/-/~TZCrKvv3sbJwM_xP/current-state/. The complete dataset used to generate Figure 2-3 and associated supplements, can be downloaded and explored via interactive dashboards via our KNIME hub workflow at https://hub.knime.com/-/spaces/-/~EcnvMwYtqylu2reV/current-state/. ACFM raw image datasets will be released on BioImage Archive.

## Acknowledgements

This work was supported by National Institutes of Health grant R21AI149275 to KKH. We thank the University of Georgia SporoCore and New York University Insectary for providing *P. berghei*-infected mosquitos. We thank William Sausman, Francisco Medrano, Beatriz Morales-Hernandez and Daniel Ferguson for assistance with live luciferase assays.

