## Supplementary Figures for "Translation inhibition efficacy does not determine the *Plasmodium berghei* liver stage antiplasmodial efficacy of protein synthesis inhibitors"

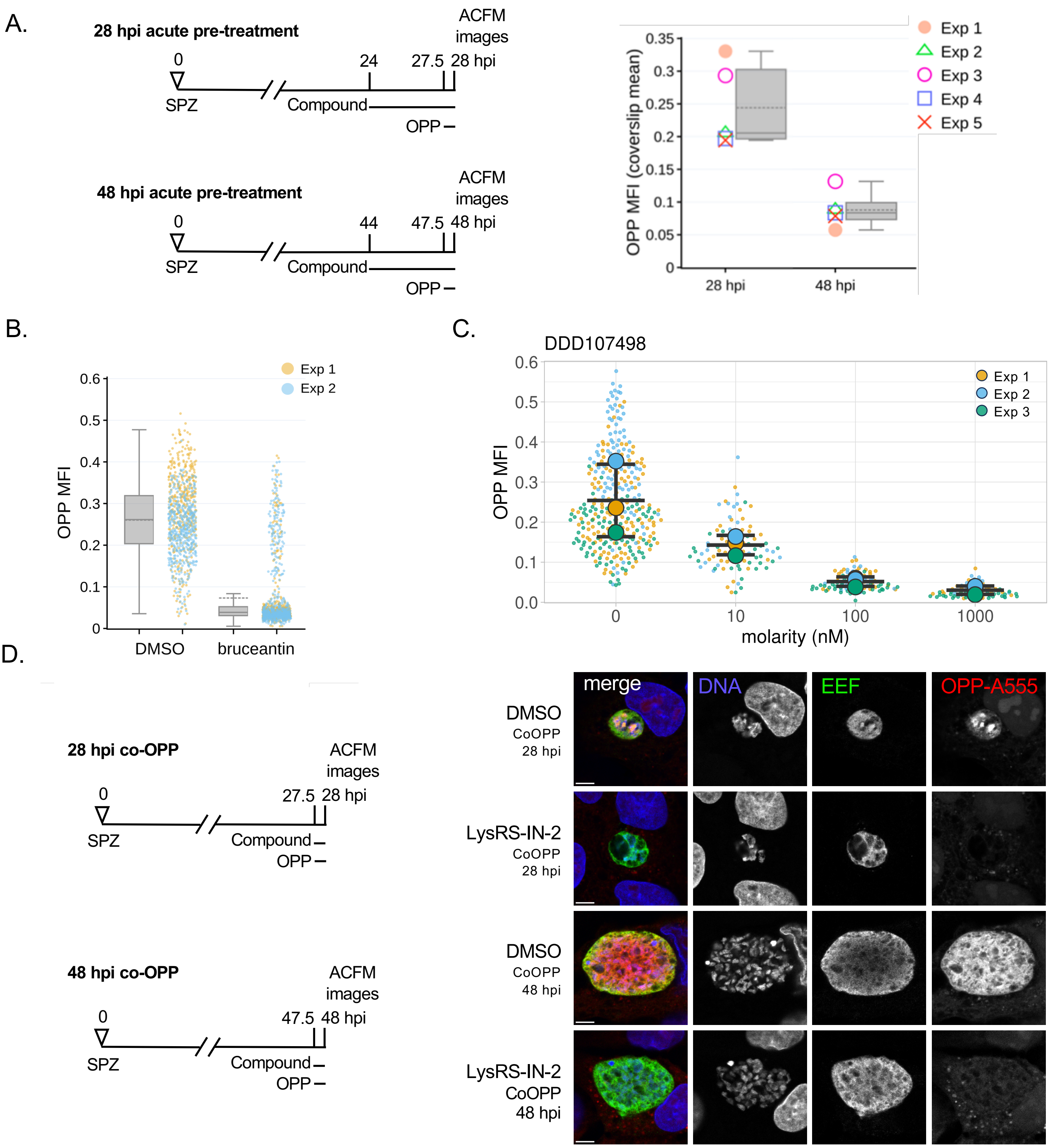


**Figure S1. Translation intensity during *P. berghei* liver stage schizogony and translation inhibitor characterization.** (A) Mean fluorescence intensity (MFI) of the nascent proteome in experimentally matched liver stage parasite populations at 28 and 48 hpi after 4 h DMSO treatment, as detailed in schematics; n=5 independent experiments, p=0.001 (two-tailed, unpaired t-test). (B) Single parasite translational intensity (OPP-MFI) at 28 hpi following a 4 hr acute pre-treatment (as in schematic in A) with 3.7 nM bruceantin or DMSO control. Boxplots show combined data from two independent experiments, with dotted lines reporting treatment means, while single points represent individual EEFs, colored by experiment. (C) SuperPlots of single parasite translational intensity (OPP-MFI) at 28 hpi following a 4 hr acute pre-treatment (as in schematic in A) with DDD107498 or DMSO control. Single points show individual EEFs, colored by independent experiment; experiment means are represented by large circles, and bars represent mean and standard deviation. This raw data is extracted from the 5-point dose response dataset published in normalized form in Fig. 3D of doi.org/10.1128/msphere.00544-23. (D) Representative confocal images of competitive LS translation inhibition by 100 µM LysRS-IN-2 vs. DMSO control, as detailed in experiment schematics. Merged images are pseudocolored as indicated with EEFs immunolabeled with α-*Pb*HSP70, OPP-A555 labeling the nascent proteome, and DNA stained with Hoechst; scale bars = 5 µm.

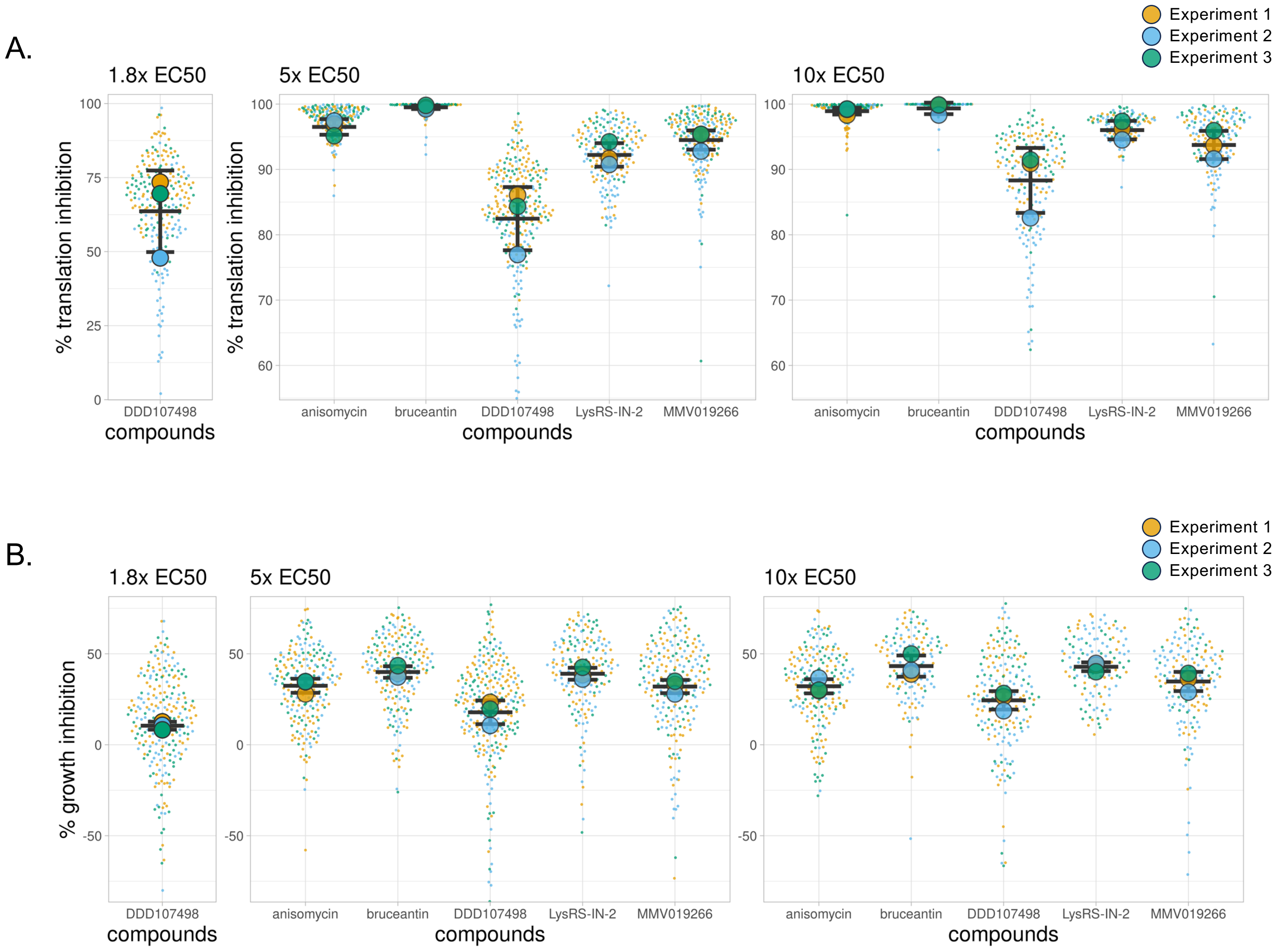


**Figure S2. Per-experiment data composition for Fig. 2.** The % translation inhibition and % growth inhibition at 28 hpi following a 4 hr acute pretreatment with individual EEFs shown as single points and experiment means are represented by large circles, color coded by independent experiment (n=3). Bars represent mean and standard deviation of the 3 experiments. Plots truncated identically to figures 2A and 2B. The full dataset can be explored via interactive dashboards in in our KNIME hub workflow: https://hub.knime.com/-/spaces/-/~EcnvMwYtqylu2reV/current-state/ .

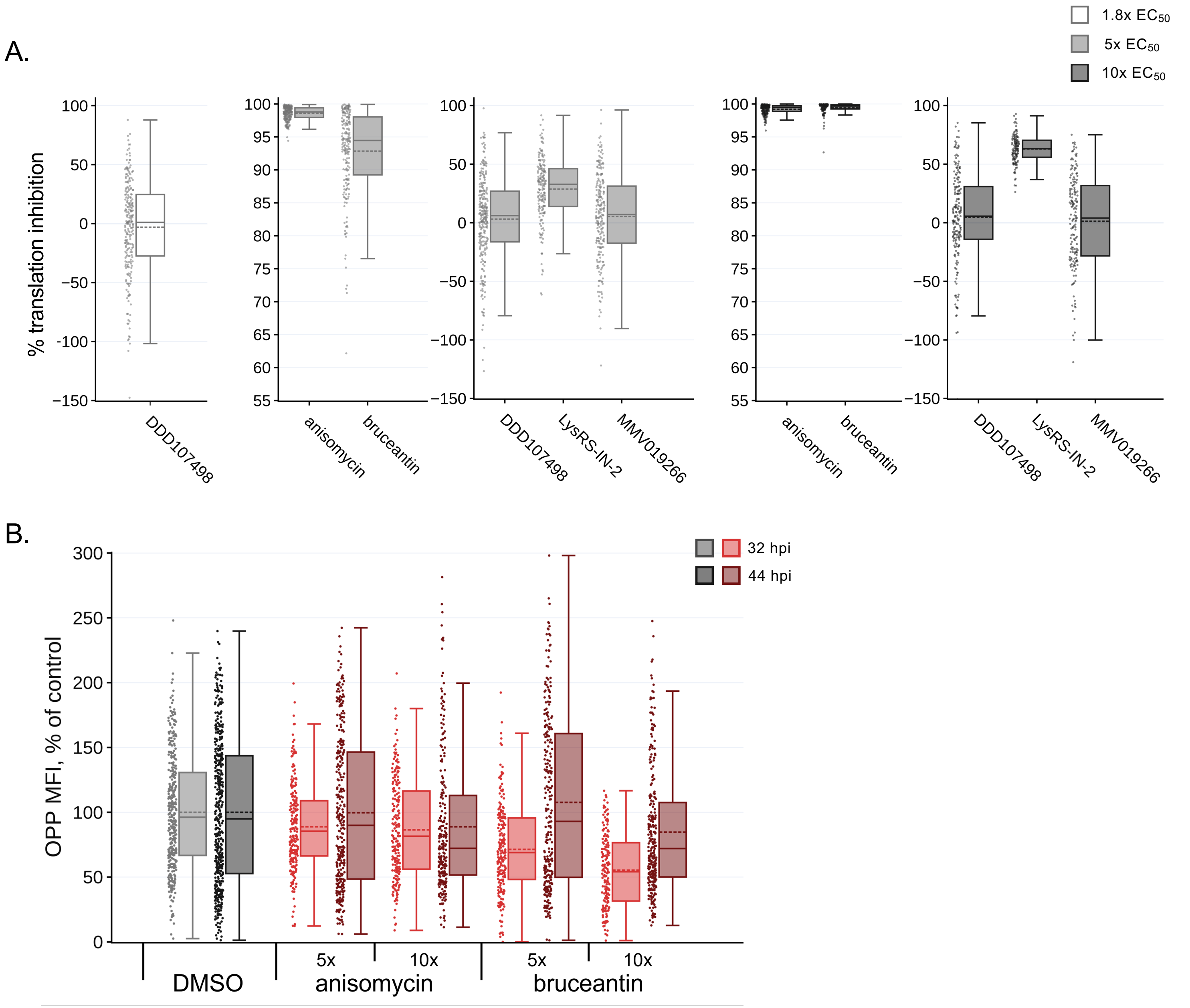


**Figure S3. Translation inhibition efficacy and recovery in HepG2 cells.** Quantification of translation in in-image HepG2 cells, corresponding to individual EEFs quantified in Figures 2 and 3, following acute pre-treatment at 28 hpi (A) and two recovery timepoints (B) for the two pan-eukaryotic active inhibitors. Individual points represent in-image HepG2 translation intensity normalized to the DMSO control mean. N=3 independent experiments. The full dataset can be explored via interactive dashboards in in our KNIME hub workflow: https://hub.knime.com/-/spaces/-/~EcnvMwYtqylu2reV/current-state/ .

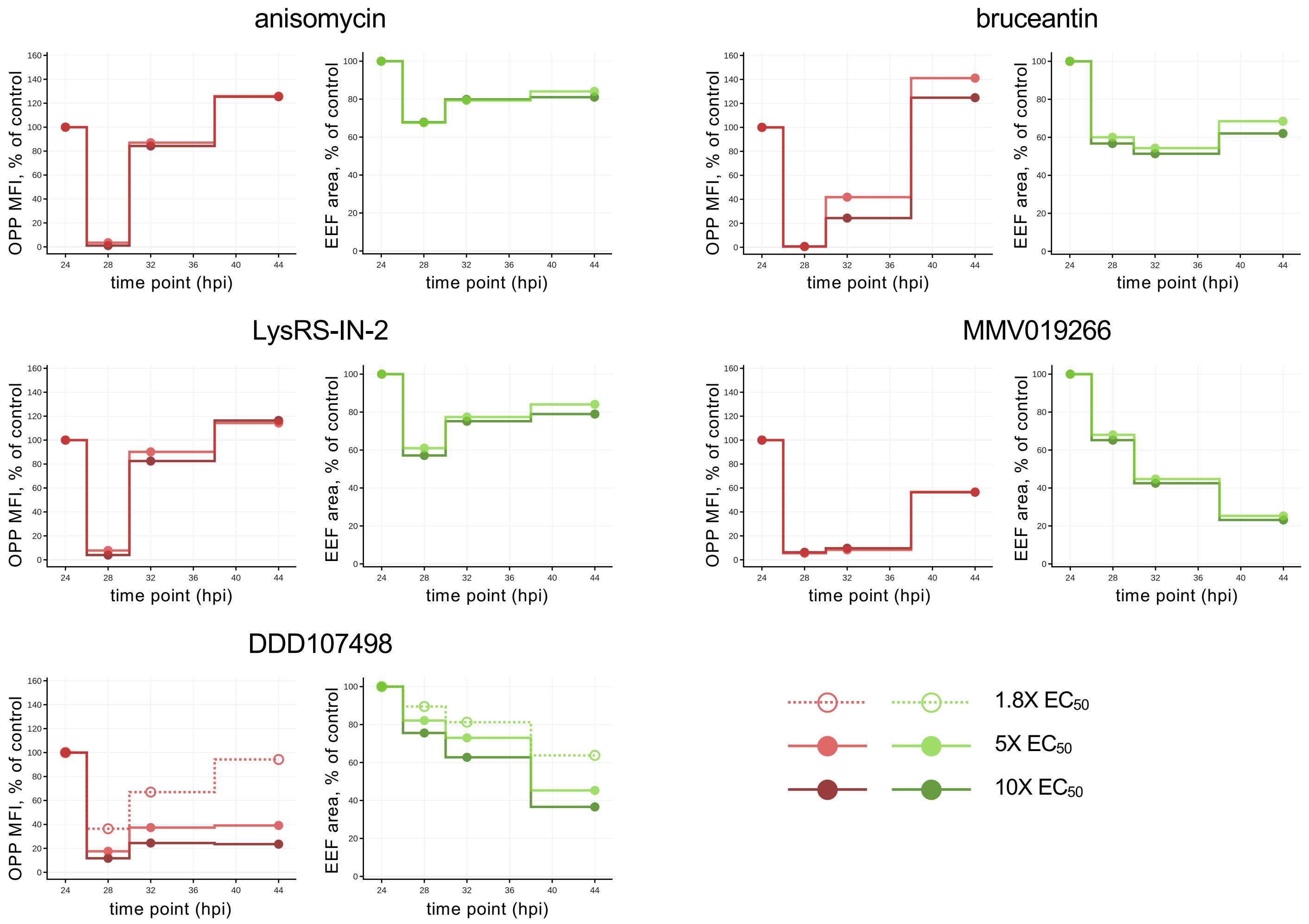


**Figure S4. Comparing translational and growth response to acute translation inhibitor treatment and washout in *P. berghei* liver stages.** Simplified line plots facilitate the comparison of effects induced by the five test compounds. OPP MFI, representing parasite translational intensity, and parasite size are normalized to the mean of timepoint matched DMSO controls. Each data point is the mean of 3 independent experiments.

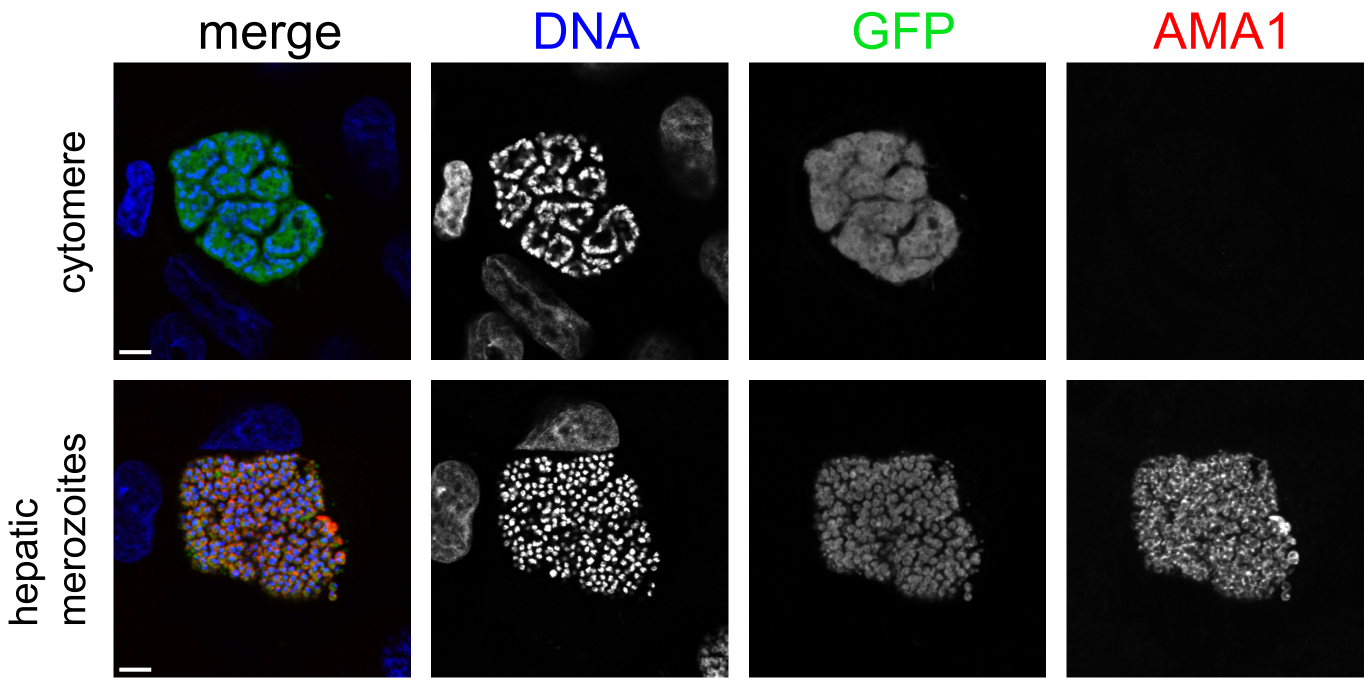


**Figure S5. AMA1 protein is present in *P. berghei* hepatic merozoites, but not in cytomere-forming late LS parasites.** Representative ACFM images of *P. berghei* EEFs at cytomere stage (top) and with hepatic merozoites (bottom) at 60 hpi. Merged images are pseudo-colored as indicated with AMA1 immunolabeled (anti-AMA1 28G2), parasite cytoplasm marked by GFP, and DNA stained with Hoechst. Scale bar = 5 µm.

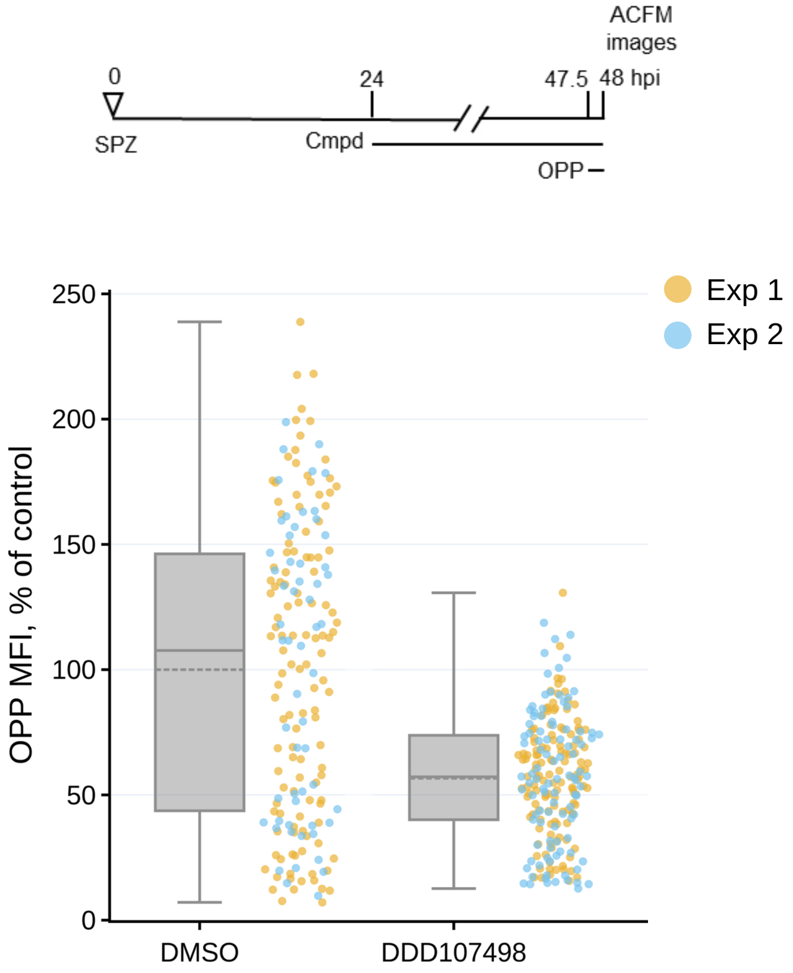


**Figure S6. DDD107498-induced translation inhibition after prolonged treatment.** Single parasite translation quantified at 48 hpi following 24 h of treatment with 20 nM DDD107498 and DMSO controls. Single data points are individual EEFs, normalized to DMSO controls from n=2 independent experiments as color coded in legend. Boxplots show combined data from both experiments, with dotted lines reporting treatment means. P <0.0001 in an unpaired, two-tailed t-test.

A.

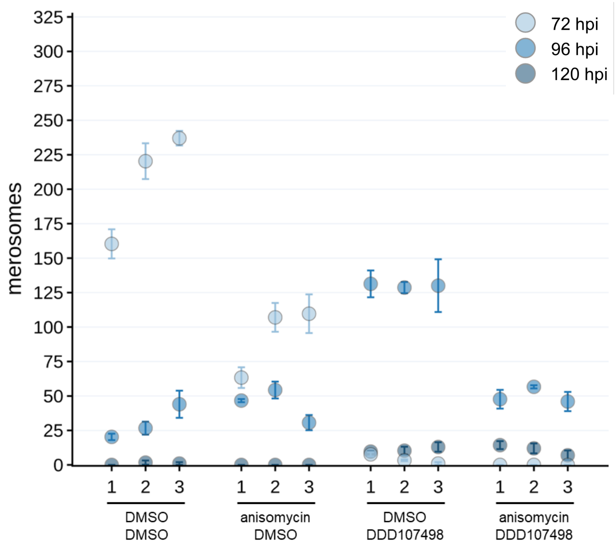


B.

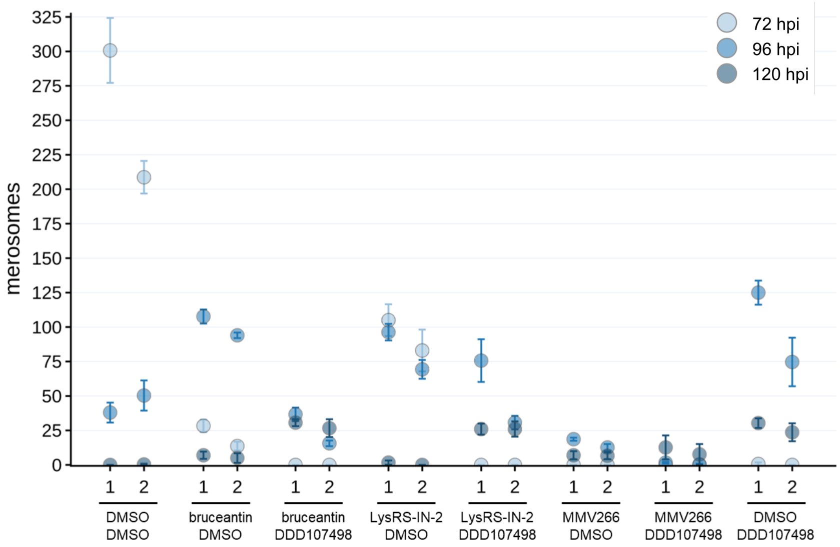


**Figure S7. DDD107498 exerts antiplasmodial effects on translationally arrested LS schizonts in each experimental replicate.** Detached cells/merosomes collected and counted in experiment-matched wells at 72, 96, and 120 hpi, with A) corresponding to Figure 6E and B) corresponding to Figure 6H. Individual data points represent mean merosome count from triplicate wells in each independent experiment (on X axis), with error bars representing standard deviations.
